# Comparative genetic architectures of schizophrenia in East Asian and European populations

**DOI:** 10.1101/445874

**Authors:** Max Lam, Chia-Yen Chen, Zhiqiang Li, Alicia R. Martin, Julien Bryois, Xixian Ma, Helena Gaspar, Masashi Ikeda, Beben Benyamin, Brielin C. Brown, Ruize Liu, Wei Zhou, Lili Guan, Yoichiro Kamatani, Sung-Wan Kim, Michiaki Kubo, Agung Kusumawardhani, Chih-Min Liu, Hong Ma, Sathish Periyasamy, Atsushi Takahashi, Qiang Wang, Zhida Xu, Hao Yu, Feng Zhu, Psychiatric Genomics Consortium - Schizophrenia Working Group, Indonesia Schizophrenia Consortium, Genetic REsearch on schizophreniA neTwork-China and Netherland (GREAT-CN), Wei J. Chen, Stephen Faraone, Stephen J. Glatt, Lin He, Steven E. Hyman, Hai-Gwo Hwu, Tao Li, Steven McCarroll, Benjamin M. Neale, Pamela Sklar, Dieter Wildenauer, Xin Yu, Dai Zhang, Bryan Mowry, Jimmy Lee, Peter Holmans, Shuhua Xu, Patrick F. Sullivan, Stephan Ripke, Michael O’Donovan, Mark J. Daly, Shengying Qin, Pak Sham, Nakao Iwata, Kyung S. Hong, Sibylle G. Schwab, Weihua Yue, Ming Tsuang, Jianjun Liu, Xiancang Ma, René S. Kahn, Yongyong Shi, Hailiang Huang

**Affiliations:** Research Division, Institute of Mental Health, Singapore, Singapore, Singapore, 539747, Singapore.; Human Genetics 2, Genome Institute of Singapore, Singapore, Singapore, 138672, Singapore.; Analytic and Translational Genetics Unit, Massachusetts General Hospital, Boston, MA, 02114, USA.; Department of Medicine, Harvard Medical School, Boston, MA, 2115, USA.; Stanley Center for Psychiatric Research, Broad Institute of Harvard and MIT, Cambridge, MA, 02142, USA.; The Biomedical Sciences Institute of Qingdao University, Qingdao Branch of SJTU Bio-X Institutes & the Affiliated Hospital of Qingdao University, Qingdao, Qingdao, 266003, China.; Bio-X Institutes, Key Laboratory for the Genetics of Developmental and Neuropsychiatric Disorders (Ministry of Education) and the Collaborative Innovation Center for Brain Science, Shanghai Jiao Tong University, Shanghai, Shanghai, 200030, China.; Department of Medical Epidemiology and Biostatistics (MEB), C8, Karolinska Instituet, Stockolm, Stockolm, SE-171, Sweden.; Chinese Academy of Sciences (CAS) Key Laboratory of Computational Biology, Max Planck Independent Research Group on Population Genomics, CAS-MPG Partner Institute for Computational Biology (PICB), Shanghai Institutes for Biological Sciences, CAS, Shanghai 200031, China, Shanghai, Shanghai, 200031, China.; University of Chinese Academy of Sciences, Beijing, Beijing, 100049, China.; Social Genetic & Developmental Psychiatry, King’s College, London, London, London, SE5 8AF, UK.; Department of Psychiatry, Fujita Health University School of Medicine, Nagoya, Aichi, 470-1192, Japan.; School of Health Sciences, University of South Australia, Adelaide, South Australia, 5000, Australia.; Institute for Molecular Bioscience, The University of Queensland, Queensland, Queensland, 4072, Australia.; Verily Life Sciences.; Shanghai Key Laboratory of Psychotic Disorders, Shanghai Mental Health Center, Shanghai Jiao Tong University School of Medicine, Shanghai, China.; National Clinical Research Center for Mental Disorders & Key Laboratory of Mental Health, Ministry of Health (Peking University), Peking University Sixth Hospital (Institute of Mental Health), Beijing, Beijing, 100191, China.; Center for Genomic Medicine, Kyoto University Graduate School of Medicine, Kyoto, Kyoto, 606-8507, Japan.; Laboratory for Statistical Analysis, RIKEN Center for Integrative Medical Sciences, Yokohama, Kanagawa, 230-0045, Japan.; Department of Psychiatry, Chonnam National University Medical School, Gwangju, South Korea, 61469, South Korea.; RIKEN Center for Integrative Medical Sciences, Yokohama, Kanagawa, 230-0045, Japan.; Psychiatry Department, University of Indonesia - Cipto Mangunkusumo National General Hospital, Jakarta Pusat, DKI Jakarta, 10430, Indonesia.; Department of Psychiatry, National Taiwan University Hospital, Taipei, Taiwan, 100, Taiwan.; Department of Psychiatry, National Taiwan University College of Medicine, Taipei, Taiwan, 100, Taiwan.; Queensland Brain Institute, The University of Queensland, Queensland, Queensland, QLD 4072, Australia.; Queensland Center for Mental Health Research, the University of Queensland Australia, Wacol, Queensland, QLD 4076, Australia.; Department of Genomic Medicine, Research Institute, National Cerebral and Cardiovascular Center, Suita, Osaka, 565-8565, Japan.; Psychiatric Laboratory, Huaxi Brain Research Center, State Key Laboratory of Biotherapy, Mental Health Center, West China Hospital of Sichuan University, Chengdu, Sichuan, 610041, China.; Department of Psychiatry, University Medical Center Utrecht, Utrecht, Utrecht, 3584 CX, Netherlands.; Center for Translational Medicine, The First Affiliated Hospital of Xi’an Jiaotong University, Xi’an, Shaanxi, 710061, China.; Clinical Research Center for Mental Disease of Shaanxi Province, The First Affiliated Hospital of Xi’an Jiaotong University, Xi’an, Shaanxi, 710061, China.; Brain Science Research Center, The First Affiliated Hospital of Xi’an Jiaotong University, Xi’an, Shaanxi, 710061, China.; Institute of Epidemiology and Preventive Medicine, National Taiwan University, Taipei, Taiwan, 100, Taiwan.; The State University of New York, Syracuse, NY, 13210, USA.; Psychiatric Genetic Epidemiology & Neurobiology Laboratory (PsychGENe Lab), Department of Psychiatry and Behavioral Sciences, SUNY Upstate Medical University, Syracuse, NY, 13210, USA.; Shanghai Center for Women and Children’s Health, Shanghai, Shanghai, 200062, China.; Department of Stem Cell and Regenerative Biology, Harvard University, Cambridge, MA, 02138, USA.; Department of Genetics, Harvard Medical School, Boston, MA, 2115, USA.; Genetics and Genomic Sciences, Icahn School of Medicine at Mount Sinai, New York, NY, 10029, USA.; University of Western Australia, Perth, WA, WA6009, Australia.; MRC Centre for Neuropsychiatric Genetics and Genomics, Institute of Psychological Medicine and Clinical Neurosciences, School of Medicine, Cardiff University, Cardiff, Cardiff, CF24 4HQ, UK.; School of Life Science and Technology, ShanghaiTech University, Shanghai, Shanghai, 201210, China.; Collaborative Innovation Center of Genetics and Development, Shanghai, Shanghai, 200438, China.; Center for Excellence in Animal Evolution and Genetics, Chinese Academy of Sciences, Kunming, Yunnan, 650223, China.; Genetics, The University of North Carolina at Chapel Hill, Chapel Hill, North Carolina, 27599, USA.; Dept. of Psychiatry and Psychotherapy, Charité - Universitätsmedizin, Berlin, Berlin, 10117, Germany.; Collaborative Innovation Center, Jining Medical University, Jining, Shandong, 272067, China.; State Key Laboratory of Brain and Cognitive Sciences, Centre for Genomic Sciences and Department of Psychiatry, Li Ka Shing Faculty of Medicine, The University of Hong Kong, Hongkong, HK China.; Department of Psychiatry, Sungkyunkwan University School of Medicine, Samsung Medical Center, Seoul, Seoul, O6351, Korea.; Centre for Medical and Molecular Bioscience, Illawarra Health and Medical Research Institute, Faculty of Science, Medicine and Health, The University of Wollongong, Wollongong, NSW, 2522, Australia.; Psychiatry, University of California, San Diego, La Jolla, CA, 32093, USA.; Department of Psychiatry, The First Affiliated Hospital of Xi’an Jiaotong University, Xi’an, Shaanxi Province, 710061, China.; Department of Psychiatry and Behavioral Health System, Icahn School of Medicine at Mount Sinai, New York, NY, 10029, USA.; Department of Psychiatry, the First Teaching Hospital of Xinjiang Medical University, Urumqi, Xinjiang, 830054, China

## Abstract

Schizophrenia is a severe psychiatric disorder with a lifetime risk of about 1% world-wide. Most large schizophrenia genetic studies have studied people of primarily European ancestry, potentially missing important biological insights. Here we present a study of East Asian participants (22,778 schizophrenia cases and 35,362 controls), identifying 21 genome-wide significant schizophrenia associations in 19 genetic loci. Over the genome, the common genetic variants that confer risk for schizophrenia have highly similar effects in those of East Asian and European ancestry (r_g_=0.98), indicating for the first time that the genetic basis of schizophrenia and its biology are broadly shared across these world populations. A fixed-effect meta-analysis including individuals from East Asian and European ancestries revealed 208 genome-wide significant schizophrenia associations in 176 genetic loci (53 novel). Trans-ancestry fine-mapping more precisely isolated schizophrenia causal alleles in 70% of these loci. Despite consistent genetic effects across populations, polygenic risk models trained in one population have reduced performance in the other, highlighting the importance of including all major ancestral groups with sufficient sample size to ensure the findings have maximum relevance for all populations.

---

Schizophrenia is an often disabling psychiatric disorder which occurs worldwide with a lifetime risk of about 1%^1^. It is well-established that genetic factors contribute to susceptibility of schizophrenia. Recently, 145 genetic loci have been associated with schizophrenia in samples of primarily European ancestry^2,3^ (EUR) but this still represents the tip of the iceberg with respect to common variant liability to the disorder: the highly polygenic nature of common variation underlying this disorder predicts that there are hundreds more loci to be discovered^4^.

Most genetic studies of schizophrenia have been in EUR samples with relatively few studies in other populations^5-8^. This is a significant deficiency for multiple reasons, particularly as it greatly limits the discovery of biological clues about schizophrenia. For some causal variants, ancestry-related heterogeneity yields varying allele frequency and linkage disequilibrium (LD) patterns such that associations that can be detected in one population may not be readily detected in others. Examples include a nonsense variant in *TBC1D4* which confers muscle insulin resistance and increases risk for type 2 diabetes that is common in Greenland but is rare or absent in other populations^9^, several Asian-specific coding variants which influence blood lipids^10^, a variant highly protective against alcoholism that is common in Asian populations but very uncommon elsewhere^11^, and two loci associated with major depression^12^that are more common in the Chinese populations than EUR^12,13^ (rs12415800: 45% versus 2%, and rs35936514: 28% versus 6%).

Even if alleles have similar frequencies across populations, the effects of alleles on risk might be specific to certain populations if there are prominent but local contributions of clinical heterogeneity, gene-environment (GxE) or gene-gene (GxG) interactions. In addition, there have been debates about differences in prevalence, symptomatology, etiology, outcome, and course of illness across geographical regions^14-19^. Understanding the genetic architecture of schizophrenia across populations provides insights in whether any differences represent etiologic heterogeneity on the illness.

Finally, polygenic risk score (PRS) prediction is emerging as a useful tool for studying the effects of genetic liability, identifying more homogeneous phenotypes, and stratifying patients, but the applicability of training data from EUR studies to those of non-European ancestry has not been fully assessed, leaving us with an uncertainty as to the biological implications and utility in non-Europeans^20^.

## Schizophrenia genetic associations in the East Asian populations

To systematically examine the genetic architecture of schizophrenia in individuals of East Asian ancestry (EAS), we compiled 22,778 schizophrenia cases and 35,362 controls from 20 samples from Singapore, Japan, Indonesia, Korea, Hong Kong, Taiwan, and mainland China (Extended Data Table 1). Individual-level genotypes were available from 16 samples (Extended Data Table 1a), on which we performed quality control, imputation and association tests (Methods and Supplementary Table 1). Two samples (TAI-1 and TAI-2) were trio-based and pseudo-controls were used. Four samples made available summary statistics for 22K-31K selected variants (Methods) which had been analyzed in published studies^7,8^.

We used a two-stage study design (Extended Data Table 1a). Stage 1 included 13 samples for which we had individual genotype data (13,305 cases and 16,244 controls after quality control). Stage 2 incorporated the remaining 7 samples: full genotype data from 3 samples that arrived after the Stage 1 data freeze and summary statistics (for selected variants) from 4 samples (Extended Data Table 1). Meta-analyses across Stage 1 samples and across all EAS samples were conducted using a fixed-effect model with inverse-variance weighting. QQ plots (Extended Data Fig. 1) showed no inflation of test statistics (particularly that ancestry effects have been well controlled) with λ_gc_=1.14, λ_1000_=1.01 and LD Score regression^21^ (LDSC) intercept=1.0145±0.011 using Stage 1 samples.

Combining Stages 1 and 2, we found 21 genome-wide significant associations at 19 loci (Table 1, Fig. 1a and Supplementary Table 2), an additional 14 associations over the most recent schizophrenia genetic study of Chinese ancestry^8^. Most associations were characterized by marked differences in allele frequencies between the EAS and EUR samples: for 15 of 21 loci, the index variants had a higher minor allele frequencies (MAF) in EAS than EUR. The higher allele frequency potentially confers better power to detect associations in EAS. For example, we identified a locus (Fig. 1b) with the top association (rs374528934) having strong evidence in EAS (*P* = 5 x 10^-11^) but not in EUR using the Stage 1 samples. rs374528934 has MAF of 45% in EAS but only 0.7% in EUR. No other variant in this locus is significantly associated with schizophrenia in EUR. This locus contains *CACNA2D2* (the calcium channel α2δ-2 subunit) associated with childhood epilepsy^22,23^, and to which the anticonvulsant medication gabapentin binds, suggesting a path for further therapeutic investigation^23^. This finding also adds new evidence to the calcium signaling pathway suggested to be implicated in psychiatric disorders^24,25^. The absence of the MHC association is evaluated in Discussion.

**Table 1.** Genome-wide significant loci in the East Asian populations. BP: genomic position in HG19. AL: Reference and non-reference alleles, OR: Odds-ratio, P: *P*-value.

| SNP | Chr | BP | AL | Stage 1 |  | Stage 2 |  | Combined |  |
| --- | --- | --- | --- | --- | --- | --- | --- | --- | --- |
|  |  |  |  | P | OR | P | OR | P | OR |
| rs4660761 | 1 | 44440146 | A/G | 3.6E-06 | 0.91 | 3.53E-04 | 0.92 | 5.08E-09 | 0.91 |
| rs848293 | 2 | 58382490 | A/G | 3.7E-10 | 0.90 | 3.10E-09 | 0.87 | 9.87E-18 | 0.89 |
| rs17592552 | 2 | 201176071 | T/C | 8.4E-10 | 0.86 | 2.68E-05 | 0.89 | 1.50E-13 | 0.88 |
| rs2073499 | 3 | 50374293 | A/G | 1.1E-09 | 0.89 | 2.14E-05 | 0.91 | 1.33E-13 | 0.90 |
| rs76442143 | 3 | 51043599 | T/C | 6.9E-09 | 1.14 | 1.03E-02 | 1.08 | 6.40E-10 | 1.12 |
| rs10935182 | 3 | 136137422 | A/G | 1.3E-06 | 0.90 | 1.33E-04 | 0.90 | 7.08E-10 | 0.90 |
| rs4856763 | 3 | 161831675 | A/G | 3.9E-06 | 0.92 | 8.54E-06 | 0.91 | 1.73E-10 | 0.92 |
| rs13096176 | 3 | 180752138 | T/C | 3.1E-07 | 0.88 | 2.21E-03 | 0.90 | 3.35E-09 | 0.89 |
| rs6832165 | 4 | 24270210 | C/G | 3.7E-08 | 1.12 | 3.70E-01 | 1.08 | 2.79E-08 | 1.12 |
| rs13142920 | 4 | 176728614 | A/C | 9.5E-05 | 0.93 | 5.85E-06 | 0.89 | 4.85E-09 | 0.92 |
| rs4479913 | 6 | 165075210 | A/G | 3.6E-07 | 1.13 | 9.98E-05 | 1.12 | 1.53E-10 | 1.12 |
| rs320696 | 7 | 137047137 | A/C | 5.5E-08 | 0.90 | 1.07E-02 | 0.93 | 2.81E-09 | 0.91 |
| rs11986274 | 8 | 38259481 | T/C | 5.1E-04 | 1.07 | 2.73E-06 | 1.11 | 1.44E-08 | 1.08 |
| rs2612614 | 8 | 65310836 | A/G | 2.2E-08 | 1.14 | 4.51E-02 | 1.06 | 1.62E-08 | 1.11 |
| rs4147157 | 10 | 104536360 | A/G | 6.6E-10 | 0.90 | 3.87E-07 | 0.89 | 1.32E-15 | 0.89 |
| rs10861879 | 12 | 108609634 | A/G | 4.8E-07 | 1.09 | 5.00E-03 | 1.07 | 1.18E-08 | 1.08 |
| rs1984658 | 12 | 123483426 | A/G | 5.1E-11 | 0.89 | 2.14E-04 | 0.92 | 8.62E-14 | 0.90 |
| rs9567393 | 13 | 32763757 | A/G | 3.5E-08 | 1.11 | 4.37E-03 | 1.07 | 1.13E-09 | 1.09 |
| rs9890128 | 17 | 1273646 | T/C | 3.5E-08 | 0.90 | 2.44E-02 | 0.91 | 2.61E-09 | 0.90 |
| rs11665111 | 18 | 77622996 | T/C | 5.2E-06 | 1.08 | 6.89E-04 | 1.09 | 1.46E-08 | 1.09 |
| rs55642704 | 18 | 77688124 | T/C | 1.1E-06 | 1.09 | 7.11E-06 | 1.10 | 3.76E-11 | 1.09 |

**Figure 1.**
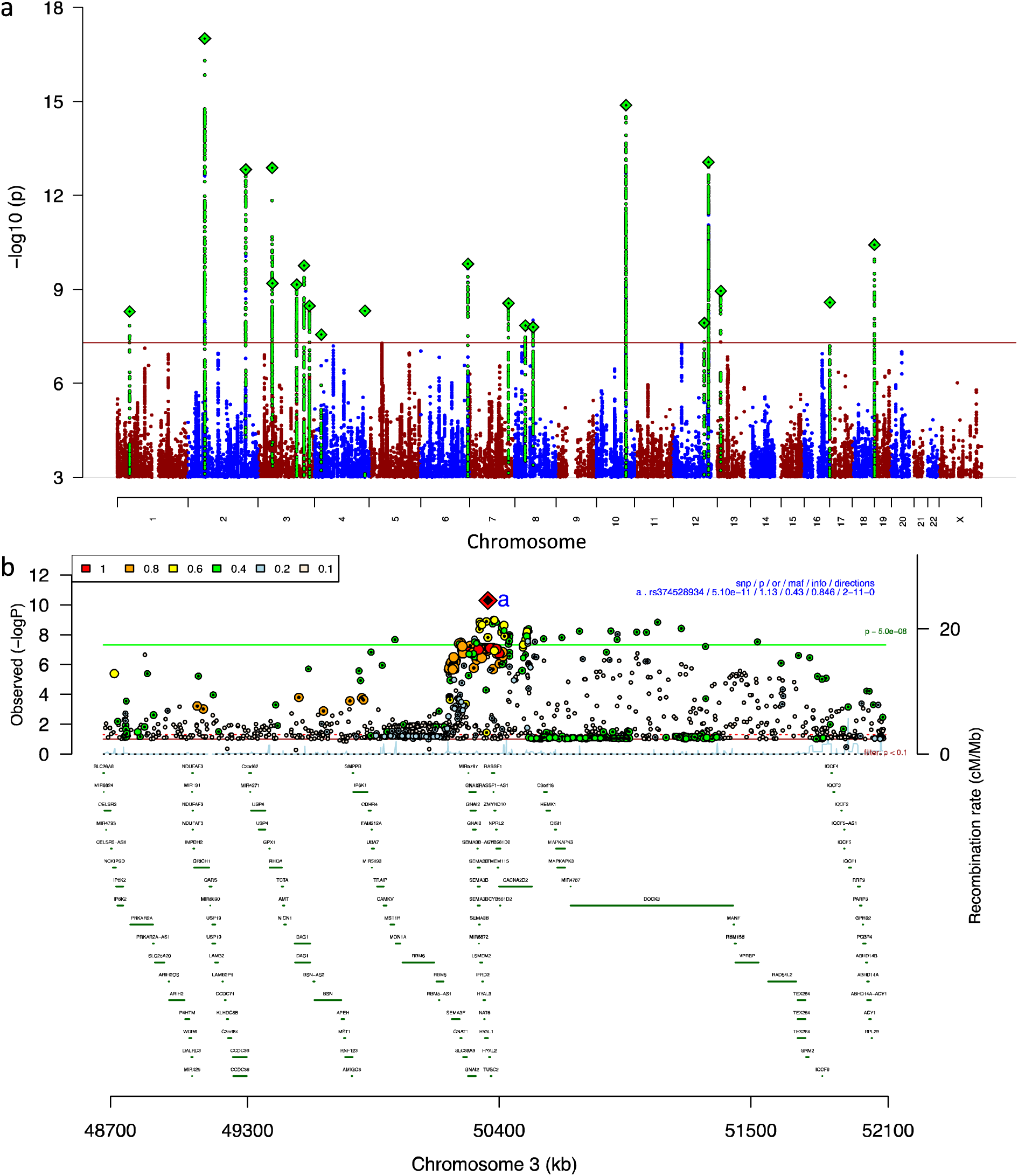
Genetic associations in East Asian populations. Horizontal line indicates the genome-wide significance threshold. **a,** Manhattan plot for schizophrenia genetic associations using East Asian samples (Stages 1 and 2). **b,** Regional association plot for a locus associated with schizophrenia using EAS Stage 1 samples.

## Genetic effects are consistent across populations

While it is assumed that biological pathways underlying complex human disorders are generally consistent across populations, genetic heterogeneity has been observed. For example, rs4246905, a variant in the *TNFSF15-TNFSF8* locus, has a much larger protective effect for Crohn’s disease in EAS than EUR (95% confidence interval of odds ratio: 0.52-0.64 vs 0.85-0. 89)^26^. For causal variants, heterogeneity of genetic effect across populations could arise from clinical heterogeneity, differences in pathophysiology, exposures to different environmental factors (GxE interaction), or interaction with other genetic factors (GxG interaction) that act non-additively with risk alleles. This large EAS sample allowed us, for the first time, to explore the heterogeneity of genetic effects influencing liability to schizophrenia across two major world populations.

Using LDSC^21^, we found the SNP-heritability of schizophrenia is very similar in EAS (0.23±0.03) and EUR (0.24±0.02) (Methods and Extended Data Fig. 2a). We also found that the common-variant genetic correlation for schizophrenia between EAS and EUR was indistinguishable from 1 (r_g_=0.98±0.03) (using POPCORN^27^, a method designed for cross-ancestry comparisons). This finding indicates that the common variant genetic architecture of schizophrenia is basically identical across EAS and EUR.

Genetic correlations between schizophrenia and 11 other psychiatric disorders and behavior traits also showed no significant differences when estimated within EUR and across EAS-EUR (Extended Data Fig.2b). In agreement with recent reports^28-31^, we observed significant positive genetic correlations for schizophrenia with bipolar disorder, major depressive disorder, anorexia nervosa, neuroticism, autism spectrum disorder, and educational attainment. We observed significant negative correlations with general intelligence, fluid intelligence score, prospective memory, and subjective well-being.

We used partitioned LDSC^21^ to look for heritability enrichment in diverse functional genomic annotations defined and used in previous publications^32,33^ (Methods and Extended Data Figure 2c,d). Using EAS Stage 1 samples, we observed significant enrichment (after Bonferroni correction) in regions conserved across 29 mammals (Conserved LindbladToh^34^). No other annotations were significantly enriched, and there were no significant differences between EUR-only and EAS-only enrichments (*P*=0.16, two-sided paired t test).

We identified gene-sets that are enriched for schizophrenia genetic associations using MAGMA^35^ and gene-set definitions from a recent schizophrenia exome sequencing study^36^ (Methods). Despite large differences in sample size and genetic background, the gene-sets implicated in EAS and EUR samples were highly consistent: we observed no significant differences between gene-set ranks using the EAS samples from the ranks using EUR samples (*P* = 0.72, Wilcoxon test). In addition, 9 of the top 10 gene-sets identified using the EAS samples are also among the top 10 gene-sets identified using EUR samples (Extended Data Figure 3).

A study of EUR individuals suggested that common schizophrenia alleles are under strong background selection^3^. We performed two analyses and found that the natural selection signatures, including positive and background selections, are consistent in schizophrenia-associated loci across EAS and EUR populations. First, we compared the signatures in the top 100 associated loci in EAS to those in EUR. Among the selection signatures we calculated (Methods), none showed a significant difference across populations (Extended Data Figure 4a, *P* > 0.05 for all panels, two-sided t test). We next asked whether the population differentiation drives schizophrenia variants to have different effect in different populations. Using 295 autosomal variants that are genome-wide significant in EAS, EUR or EAS-EUR combined samples, we did not observe a correlation (*R*^2^=0.003, Extended Data Figure 4b) between the population differentiation (measured by *F_st_)* and the heterogeneity of effect size (measured by log_10_P-value from the heterogeneity test across EAS and EUR).

We compared the effect size estimates for schizophrenia associations in EAS versus those in EUR. A precise comparison requires disease-causal variants and equivalent case and control ascertainment schemes to avoid heterogeneity driven by differences in LD and heterogeneity due to differences in cases and in controls. As we do not know the causal alleles at the associated loci, we used the most significantly associated variants in EAS that are in LD (*R*^2^>0.8) with the most significantly associated variants in EUR at each locus as an approximation. We also restricted the comparison to variants that have *P*<10^-10^ in EUR and MAF > 10% in EAS as the estimates of the effect sizes for relatively common alleles that substantially surpass genome-wide significance are least subject to inflationary bias in the discovery set. None of the 21 associations that met these criteria showed significant differences in the direction of effect (Fig. 2a) and moreover, the magnitude of the effect size was consistent across the two populations with a modest bias from the winner’s curse in the discovery (EUR) samples (slope=0.67±0.09).

**Figure 2.**
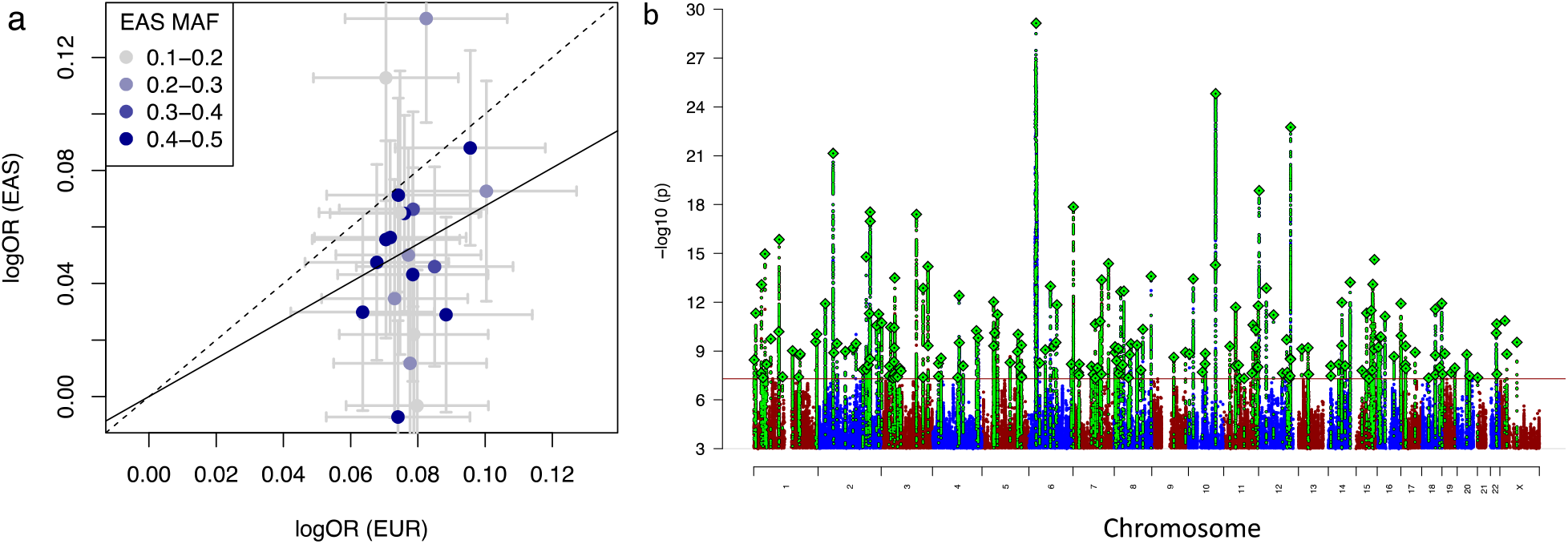
Schizophrenia associations in EUR and EAS samples. **a,** Log odds ratio of top schizophrenia associations estimated in EUR and EAS samples. Error bars indicate 95% confidence interval. Dashed line indicates the diagonal line, and the solid line indicates the regression line with intercept at 0. **b,** Manhattan plot for the schizophrenia genetic associations from the EAS (Stages 1 and 2) + EUR meta-analysis.

## Schizophrenia genetic associations from the meta-analysis of EAS and EUR

As the genetic effects observed in EAS are largely consistent with those observed in EUR, we performed a meta-analysis including the EUR and EAS samples (Stages 1 and 2) using a fixed-effect model with inverse-variance weighting^37^. The EUR samples in this analysis (56,418 cases and 78,818 controls) included all samples of EUR ancestry from the previous publication^2^ with the exclusion of three samples of EAS ancestry and the deCODE samples (1,513 cases and 66,236 controls) which only had summary statistics for selected variants. The three EAS samples (IMH-1, HNK-1 and JPN-1) excluded from EUR samples were included in our EAS Stage 1.

We identified 208 independent (both in EAS and EUR) variants associated with schizophrenia across 176 genetic loci (Fig. 2b and Supplementary Tables 3 and 4), among which 53 loci were novel (not reported in ref 2,3,7,8). Of the 108 schizophrenia-associated loci reported in the previous EUR study^2^, 89 remained significant in this study (Supplementary Table 5). As suggested by Pañrdias *et al.^3^*, this reflects an expected over-estimation of the effect sizes due to the winner’s curse in the previous study, but does not mean the 19 loci not significant in this study were false-positives in the previous study. In addition, deCODE samples were not included in this analysis.

## Population diversity improves fine-mapping

Due to LD, disease-associated loci from genome-wide association studies usually implicate genomic regions containing many associated variants. A number of approaches allow for the associated variants to be refined to a smaller set of the most plausible (or credible) candidate causal variants^38-41^. Loci implicated in psychiatric disorders usually have small effect sizes and as a result, have generally poor performance using such approaches^2,3^.

Diversity in genetic background across populations can be used to improve fine-mapping resolution^42^. Here we demonstrate that resolution can be improved by exploiting differences in the patterns of LD between causal (directly associated) and LD (indirectly) associated variants. Based on the premise that genetic effects are highly consistent across populations, the causal variants will have consistent effects across populations whereas non-causal variants can have inconsistent effects due to population-specific LD patterns. We therefore expect causal variants to have greater statistical significance and less heterogeneity in trans-ancestry meta-analysis compared to other alleles that are indirectly associated via LD (Extended Data Figure 5). Using a new algorithm based on this presumption (Methods), we fine-mapped 133 schizophrenia associations that reached genome-wide significance in the EUR and EAS (Stage 1) combined meta-analysis (Supplementary Table 6). Stage 2 EAS samples were excluded because not all had full genome coverage, which confounds the fine-mapping outcome (Methods).

Results from this EAS-EUR trans-ancestry approach improved upon those using only EUR, with 93 loci mapped to a smaller number of candidate causal variants. For example, a locus on chromosome 1 (238.8-239.4 Mb) which initially contained 7 potentially causal variants based on a published fine-mapping method^38^ and EUR samples was resolved to a single variant, rs11587347, with 97.6% probability (Fig. 3a). This variant showed strong association in both populations, while the other 6 variants are equally associated in EUR but not in EAS (Fig. 3b, c). Over all associations, the median size of the 95% credible set, defined as the minimum list of variants that were >95% likely to contain the causal variant, dropped from 57 to 34; and the number of associations mapped to ≤5 variants increased from 8 to 15 (Fig. 3d). The number of associations mapped to a single variant with greater than 50% probability increased from 16 to 20, and median size of the genomic regions the associations mapped decreased from 277 Kb to 111 Kb.

**Figure 3.**
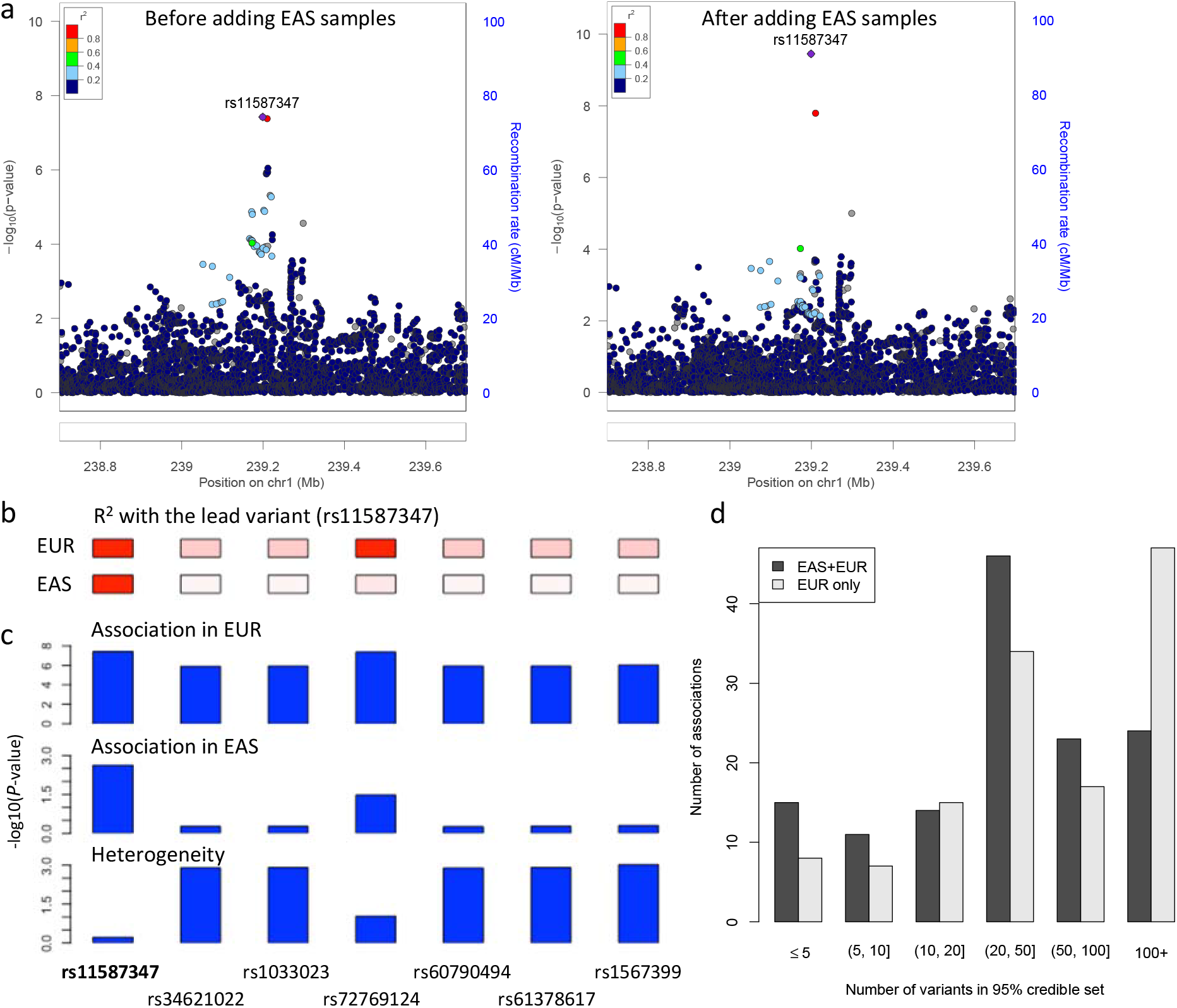
Trans-ethnicity fine-mapping maps improves resolution. **a,** an association was mapped to a single variant (rs11587347) after adding EAS samples and using the transancestry fine-mapping approach. Regional association plots were generated using http://locuszoom.org/ and LD from 1000 Genomes Project Phase 3 EUR subjects. **b,** LD with the lead variant (rs11587347). Red: perfect LD (R^2^=1); white: no LD (R^2^=0). **c,** The lead variant (rs11587347) has strong association significance in both populations and low heterogeneity across populations. **d,** Number of variants in the 95% credible set using the trans-ancestry (EAS+EUR) and publish fine-mapping approaches (EUR only).

Two schizophrenia associations were fine-mapped to coding variants including *SLC39A8* (A391T) with 44.8% probability, and *WSCD2* (T226I) with 14.8% probability. The *SLC39A8* A391T variant causes deficiency in manganese homeostasis^43^ and glycosylation^44^, and is associated with Crohn’s disease^45^, human gut microbiome composition^45^, hypertension^46^ and intelligence^47^. In addition, using a similar strategy as in Huang *et al*^38^, we found a schizophrenia association (mapped to rs1700006 with 16.1% causal probability) implicating a conserved transcription factor binding site (MEF2), which is 14 kb downstream of the nicotinic receptor subunits *CHRNA3* and *CHRNB4.* Finally, we searched for but did not find any associations that implicate constrained nucleotides near exon splicing junctions^48^.

## Transferability of genetics across populations

We compared the variance explained across EAS and EUR for genome-wide significant loci, approximated as 2*f* 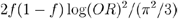 (ref 49), which explain >0.05% of the variance in either ancestry (Extended Data Fig. 6). While these variants most often have the same effect across populations, their allele frequencies can differ. Variance explained, combining the effect size (OR) and prevalence of the risk allele 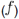, can be regarded as an approximate measure of the importance of a causal variant in a population. We found that most of the difference in variance explained is driven by allele frequency differences. One of the implications of this observation, as suggested in recent studies^20,50,51^, is that even if the risk alleles and effect sizes are primarily shared across populations, the disease predictive power of individual alleles, and of composite measures of those risk alleles such as PRS, may not be equivalent across populations.

Here we evaluate this empirically. We assessed how much variation in schizophrenia risk can be explained in EAS using both EAS Stage 1 and EUR training data. Using a standard clumping approach, we first computed PRS using a leave-one-out meta-analysis approach with EAS summary statistics (Methods), which explained ~3% of schizophrenia risk using genome-wide variants on the liability scale (*R*^2^ = 0.029 at *P*=0.5). In contrast, when EUR summary statistics were used to calculate PRS in the EAS samples, a maximum of only ~2% of schizophrenia risk was explained (*R*^2^ = 0.022 at *P*=0.1) despite a greater than 3-fold larger EUR effective sample size (Fig. 4 and Extended Data Fig. 7). The variance explained across various *P*-value thresholds provides a proxy for the signal-to-noise ratio, which differs by training population-relative to the EUR training data, variants from the EAS training data with more permissive *P*-values improve the EAS prediction accuracy. These results indicate that larger EAS studies will be needed to explain similar case/control variance as currently explained in EUR individuals. Further, although individual loci typically have the same direction and similar magnitude across populations, aggregating variants that differentially tag causal loci across populations for genetic risk prediction results in considerable variability in prediction accuracy.

**Figure 4.**
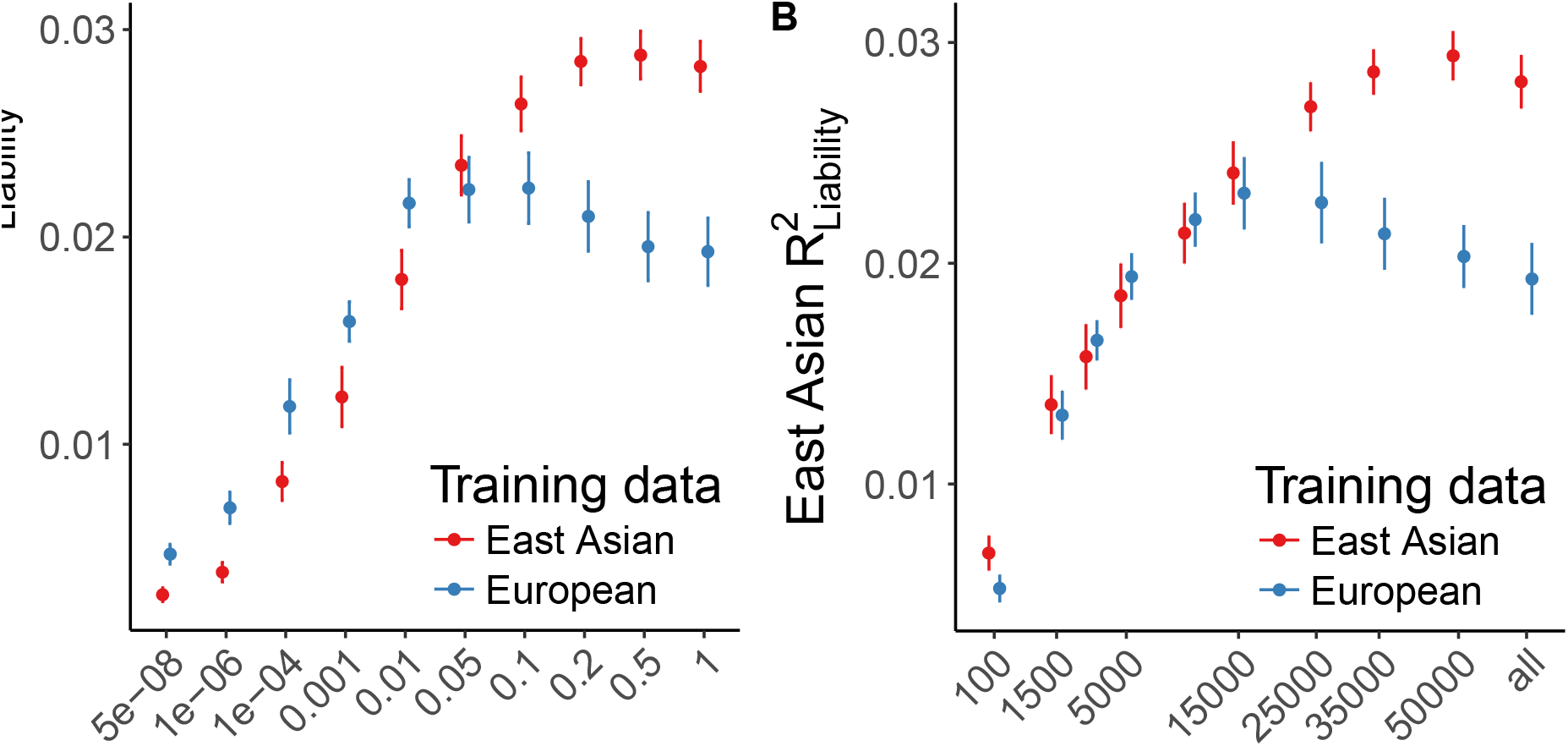
Genetic risk prediction accuracy in EAS from EAS or EUR training data. Polygenic risk scores were computed with GWAS summary statistics from EAS and EUR populations as training sets. EAS risk alleles and weights were computed with a leave-one-out meta-analysis approach across the 13 Stage 1 samples. Error bars indicate the 95% confidence interval. LD panel for clumping is from EUR and EAS 1000 Genomes Phase 3 samples. **a,** Case/control variance explained in EAS samples by variants from EAS and EUR training data with a *P*-value more significant than the threshold. **b,** Case/control variance explained by the *n* most significant independent variants.

## DISCUSSION

To date, most large-scale psychiatric genetics studies have been based on samples of primarily EUR ancestry^6^. To increase global coverage, we compiled the largest non-European psychiatric genetics cohort to date, and leveraged its size and diversity to provide new insights into the genetic architecture of schizophrenia. This study included all available major genotyped schizophrenia samples of East Asia ancestry, and presented analyses that had never been performed with sufficient power in psychiatric genetics.

When a single population is used to identify the disease-associated loci, the discovery is skewed towards disease-associated variants that have greater allele frequency in that population (Extended Data Figure 8). When multiple populations are used, disease-associated variants are equally represented across the allele frequency spectrum in these populations (Extended Data Figure 8). This demonstrates that including global samples improves power to find disease associations for which the power varies across populations. In this study, for example, more EUR than EAS samples would be required to detect around half of the new loci, as the MAF is higher in EAS than in EUR in these loci.

For traits like body mass index and autoimmune diseases, we observed heterogeneity across populations in genetic effects^26,52^, which may point to interactions between genetic associations and environment factors and/or other genetic loci. In contrast, for schizophrenia, we did not find significant heterogeneity across EAS and EUR ancestries. Analyses in genetic heritability, genetic correlation, gene-set enrichment and natural selection signatures all converge to the same conclusion that the schizophrenia biology is substantially shared across EAS and EUR, and likely, across other major world populations. This remarkable genetic correlation (*r*_g_=0.98) across populations suggests, for the first time, that schizophrenia genetic factors operate in an obedient fashion between ethnic and cultural backgrounds, and schizophrenia across the world share the same genetic causes. Given that the mainstay epidemiological factors (migration, urbanity and substance misuse) differ across populations, this finding also suggests any specific genetic liability to schizophrenia acting via these routes is minimal.

We note that a direct comparison of the effect sizes estimated in EAS with those estimated in EUR has reduced accuracy as we do not know the exact schizophrenia causal variants. This is further complicated by inflation in effect size estimates due to the winner’s curse, which are of different magnitudes due to the sample size. Increasing the sample size, especially in those of non-European ancestries, will reduce the bias and enable a better isolation of causal variants, leading to a more precise comparison of the genetic effect size across populations.

The major histocompatibility complex (MHC) hosts the strongest schizophrenia association in EUR^53^. In this study, we did not find a significant schizophrenia association in MHC in EAS. An earlier EUR study^54^ mapped the MHC associations to a set of variants (in LD) at both distal ends of the extended MHC (lead variant: rs13194504) and the complement component 4 (*C4*). Consistent with several studies of the Chinese ancestries^7,8,55,56^, none of these associations was significant in EAS in this study. We attribute this partially to low frequencies: rs13194504 has MAF < 1% in EAS comparing with 9% in EUR, and the C4-BS allele is extremely uncommon in samples from China and Korea^57,58^. Another reason may be the EUR-specific LD. In EUR, multiple protective alleles that contribute to the MHC associations are all on the same haplotype across about 6 Mb, due to an extremely long and EUR-specific haplotype that generates LD patterns at 5-Mb scale. This is also the reason that that association signals span so many Mb of genome, and the aggregate association signal (at variants that are in partial LD to multiple signals) is stronger than the signals at the individual associations.

Two recent studies using individuals of Chinese ancestries^7,8^ reported variants in MHC significantly associated with schizophrenia (rs115070292 and rs111782145 respectively, with very weak LD with each other: *R*^2^=0.07), which are different and not in LD with the EUR MHC associations. rs115070292, from Yu *et al.^7^*, is more frequent in EAS (12%) than in EUR (2%) with *P* = 10^-9^ using 4,384 cases and 5,770 controls of Chinese ancestry. This variant was not significantly associated in our study (*P* = 0.44) even though some samples of the Chinese studies were included in the current study (BJM-1, 1,312 cases and 1,987 controls). OR estimated from these shared samples marginally differs from that estimated using all EAS samples (*P*=0.018), and this association showed marginally significant heterogeneity across all EAS samples (P=0.039). Similarly, we did not replicate the association at rs111782145 from Li *et al^8^* (*P* = 0.47) despite of the sample overlap (2,555 cases and 3,952 controls). Further investigation with more samples is needed to delineate MHC associations in EAS and Chinese.

Genetic associations usually implicate a large genomic region and thus it can be challenging to map their molecular functions. We designed a novel algorithm to leverage the population diversity to fine-map schizophrenia associations to precise sets of variants. Using this algorithm we reduced the number of candidate variants associated with schizophrenia and facilitated the functional interpretation of these associations. Our algorithm assumed that there is a single causal variant in a genetic locus associated with schizophrenia. Previous fine-mapping studies^38,59^ have confirmed that this assumption is valid for most genetic loci associated with complex disorders.

Finally, this large-scale EAS sample allowed us to empirically evaluate the congruence of the genetic basis of schizophrenia between EAS and EUR. In spite of a cross-population genetic correlation indistinguishable from 1, we found that polygenic risk models trained in one population have reduced performance in the other population due to different allele frequency distributions and LD structures. This highlights the importance of including all major ancestral groups in genomic studies both as a strategy to improve power to find disease associations and to ensure the findings have maximum relevance for all populations.

**Supplementary Information** is available in the online version of the paper.

## Acknowledgements

We thank Kenneth Kendler, John McGrath, James Walters, Douglas Levinson and Michael Owen for helpful discussions. We thank SURFsara and Digital China Health for computing infrastructure for this study. M.L. acknowledges National Medical Research Council Research Training Fellowship award (Grant No.: MH095:003/008-1014); Z.L. acknowledges the Natural Science Foundation of China (NSFC 81701321); B.B. acknowledges funding support from the National Health and Medical Research Council (NHMRC Funding No. 1084417 and 1079583); Y.K., M.K. and A.T. acknowledge Strategic Research Program for Brain Sciences (SRPBS) from Japan Agency for Medical Research and Development (AMED); part of the BioBank Japan Project from the Ministry of Education, Culture, Sports, and Technology (MEXT) of Japan; S-W.K. acknowledges Grant of the Korean Mental Health Technology R&D Project (HM15C1140); W.J.C. acknowledges Ministry of Education, Taiwan (‘Aim for the Top University Project’ to National Taiwan University, 2011-2017); Ministry of Science and Technology, Taiwan (MOST 103-2325-B-002-025); National Health Research Institutes, Taiwan (NHRI-EX104-10432PI); NIH/NHGRI grant U54HG003067; NIMH grant R01 MH085521; and NIMH grant R01 MH085560; S.J.G. acknowledges R01MH08552; H-G.H. acknowledges Ministry of Education, Taiwan (‘Aim for the Top University Project’ to National Taiwan University, 2011-2017); MOST, Taiwan (MOST 103-2325-B-002-025); NIH/NHGRI grant U54HG003067; NIMH grants R01 MH085521, R01 MH085560; and NHRI, Taiwan (NHRI-EX104-10432PI); T.L. acknowledges National NSFC Key Project (81630030 and 81130024, PI: Tao Li); National NSFC/Research Grants Council of Hong Kong Joint Research Scheme (81461168029, PI: Tao Li and Pak C Sham); and National Key Research and Development Program of the Ministry of Science and Technology of China (2016YFC0904300, PI: Tao Li); P.S. acknowledges U01MH109536; J.L. acknowledges National Medical Research Council Translational and Clinical Research Flagship Programme (Grant No.: NMRC/TCR/003/2008) and National Medical Research Council under the Centre Grant Programme (Grant No.: NMRC/CG/004/2013); S.X. acknowledges financial support from the Strategic Priority Research Program (XDB13040100) and Key Research Program of Frontier Sciences (QYZDJ-SSW-SYS009) of the Chinese Academy of Sciences (CAS), the National Natural Science Foundation of China (NSFC) grant (91731303, 31771388, and 31711530221), the National Science Fund for Distinguished Young Scholars (31525014), the Program of Shanghai Academic Research Leader (16XD1404700), the National Key Research and Development Program (2016YFC0906403); and Shanghai Municipal Science and Technology Major Project (2017SHZDZX01); P.F.S. acknowledges the PGC funding from U01 MH109528 and U01 MH1095320; S.Q. acknowledges National Key Research and Development Program of China (2016YFC0905000, 2016YFC0905002); and Shanghai Key Laboratory of Psychotic Disorders (13dz2260500); K.S.H. acknowledges Grant of the National Research Foundation of Korea (2015R1A2A2A01002699); W.Y. acknowledges National Key R&D Program of China(2016YFC1307000); National Key Technology R&D Program of China (2015BAI13B01); National NSFC (81571313, 91232305, 91432304); M.T. acknowledges R01MH085560: Expanding Rapid Ascertainment Networks of Schizophrenia Families in Taiwan; X.M. acknowledges the National NSFC Surface project (81471374; PI: XC Ma); Y.S. acknowledges National Key R&D Program of China (2016YFC0903402); NSFC (31325014, 81130022, 81421061); and the 973 Program (2015CB559100). H.H. acknowledges NIDDK K01DK114379 and funding support from Stanley Center for Psychiatric Research.

## Author contribution

Genotype quality control and principal component analyses: M.L., H.H.; Association analysis: M.L., C.C.; Heritability and genetic correlation: M.L. B.C.B; Natural selection: X.M., C.C., S.X.; Partitioned heritability: J.B.; Gene-set analysis: Z.L., M.L., G.H.; Polygenic risk score: A.R.M, C.C., R.L.; Fine-mapping: H.H.; Data acquisition, generation, quality control and analysis: IMH- 1,2: M.L., J.L., J.J.L.; HNK-1: Q.W., T.L., P.S.; JPN-1: A.T., Y.K., M.K., M.I., N.I.; BIX-1-3,5; Z.L., L.H., Y.S.; XJU-1: F.Z., X.M.; UMC-1, SIX-1: L.G., H.M., Z.X., P.S., X.Y., R.S.K.; UWA-1: B.B., A.K., D.W., S.G.S.; BJM1-4: H.Y., D.Z., W.Y.; TAI-1,2: C.L., W.J.C., S.F., S.J.G., H.G.H., S.M., B.M.N., M.T.; KOR-1: S.K., K.S.H.; BIX-4: W.Z., L.H., S.Q.; Primary drafting of the manuscript: M.L., C.C., S.S., M.O., M.J.D., H.H.; Major contribution to drafting of the manuscript: A.R.M, S.G., B.B., S.P., B.M., K.S.H., M.T., J.L., W.Y., H.G.H., J.B., S.R.; Project conception, design, supervision and implementation: H.H., Y.S., R.S.K., X.M., J.L., M.T., W.Y., S.S., K,S.H., N.I., P.S., S.Q., M.J.D., M.O., S.R., P.S., L.H., S.E.H.. All authors saw, had the opportunity to comment on, and approved the final draft.

## Author information

The study protocols were approved by the institutional review board at each center involved with recruitment. Informed consent and permission to share the data were obtained from all subjects, in compliance with the guidelines specified by the recruiting centre’s institutional review board. Samples that were recruited in mainland China were processed and analyzed in a Chinese server. The authors declare no competing interests. Correspondence and requests for materials should be addressed to S.Q., P.S., N.I., K.S.H., S.G.S., W.Y., M.T., J.J.L., X.M., R.S.K., Y.S., or H.H.

## METHODS

### Overview of samples

#### The following samples were used in this study

##### EAS samples, full-genome

genome-wide genotype data was obtained from 16 EAS samples from Singapore, Japan, Indonesia, Korea, Hong Kong, Taiwan, and mainland China (Extended Data Table 1). Two of these samples (TAI-1 and TAI-2) had parents off-spring trios, and were processed as case/pseudo-controls. DSM-IV was used for diagnosing all schizophrenia cases in these samples except for the tros (TAI-1 and TAI-2), for which DIGS was used. All samples were processed according to quality control (QC) procedures reported in ref 2, with details reported in following sections. After QC, genotypes were phased and imputed against the 1000 Genomes Project Phase 3 reference panel^6^. Principal component analysis (PCA) was conducted across samples via imputed best guess genotypes to identify and remove overlapping samples across datasets, cryptic related samples and population outliers. Eight principal components (PCs) that are associated to the case-control status were included in univariate logistic regression to control for the population stratification in each sample.

##### EAS samples, selected variants

summary statistics was obtained for a set of variants from four EAS samples (BJM-2, BJM-3, BJM-4, BIX-5) which had been analyzed in published studies^7,8^. The summary statistics included odds ratio, standard error, reference and tested alleles for variants that have *P*<10^-5^ in either Stage 1 or the meta-analysis combining Stage 1 and EUR samples. Between 22,156 and 31,626 variants were available after the exclusion of strand ambiguous^60^ variants (Supplementary Table 1).

##### EUR samples

Genotypes for EUR schizophrenia patients and controls were obtained from the Psychiatric Genomics Consortium as reported in ref 2. All samples of EUR ancestry were included in this study except for the deCODE samples (1,513 cases and 66,236 controls). We also like to note that three samples of EAS ancestry reported in this publication were not included in the EUR samples in our analysis but were included in the EAS samples (IMH-1, HNK-1 and JPN-1). The same procedures used in processing EAS samples were applied to the EUR samples.

### Quality control

Quality control procedures were carried out as part of the RICOPILI pipeline (http://sites.google.com/a/broadinstitute.org/ricopili/home) with the following steps and parameters: 1) Excluding variants with call rate below 95%; 2) Excluding subjects with call rate below 98%; 3) Excluding monomorphic variants; 4) Excluding subjects with inbred coefficient above 0.2 and below -0.2; 5) Excluding subjects with mismatch in reported gender and chromosome X computed gender; 6) Excluding variants with missing rate differences greater than 2% between cases and controls; 7) Subsequent to step 6, exclude variants with call rate below 98%; and 8) Exclude variants in violation of Hardy-Weinberg equilibrium (*P* < 10^-6^ for controls or *P* < 10^-10^ for cases). Numbers of variants or subjects removed in each step were reported in Supplementary Table 1.

### Phasing and imputation

All datasets were phased using SHAPEIT^61^ and IMPUTE2^62^ using regular steps and parameters. Additional processing for trios (TAI-1 and TAI-2) was carried out such that case/pseudo-controls were identified and imputed. All samples were imputed to the 1000 Genomes Project Phase 3 reference panel^63^ (2504 subjects, including 504 EAS subjects). Imputation procedures resulted in dosage files and best guess genotypes in PLINK^64^ binary format. The former was used for subsequent association analysis and the latter was used in the PCA and PRS analyses.

### Sample overlaps, population outliers and population stratification

We used Eigenstrat^65^ to calculate the principal components for all the samples using the best guess genotypes from imputation (Extended Data Figure 9b). We computed the identity-by-descent matrix to identify intra- and inter-dataset sample overlaps. Samples with pi-hat > 0.2 were extracted, followed by Fisher-Yates shuffle on all samples. The number of times with which each sample was related to another sample was tracked and samples that were related to more than 25 samples were removed. When deciding which samples to retain, trio were preferred, followed by cases, and thereafter a random sample for each related pair was removed, 704 individuals were removed.

To identify population outliers, k-means clustering was conducted using the first 20 PCs from PCA and covariates representing each of the 13 Stage 1 samples. Guided by results of k-means clustering and visual inspection of PCA plots, 46 individuals were identified as outliers and were excluded. Further population-level inspection was carried out by merging the 1000 Genomes Project Phase 1 reference samples with Stage 1 samples and conducting PCA (Extended Data Figure 9a). Using similar approaches reported above, no further samples were excluded as population outliers.

Eight PCs that are associated with case/control status with *P* < 0.2 were used as covariates for association analysis in each sample (PCs 1, 4, 5, 6, 8, 9, 15, and 19). QQ plots (Extended Data Figure 1) showed that the population structure has been well controlled.

### Association analysis and meta-analysis

Association analysis was carried out for each sample using PLINK^64^ and genotype dosage from imputation. Only variants having imputation INFO ≥ 0.6 and MAF ≥ 1% were included in the analysis. We performed logistic regression with PCs identified in the prior subsection as covariates to control for population stratification within each study. Fixed-effect meta-analysis^66^, weighted by inverse-variance, was then used to combine the association results across samples. Meta-analysis for European samples were conducted in the same matter. In order to find independent schizophrenia associations in both EUR and EAS populations (Supplementary Table 4), we performed LD clumping twice using the 1000 Genomes Project Phase 3 EUR and EAS reference panels respectively (with default parameters in RICOPILI).

### Chromosome X analysis

Chromosome X genotypes were processed separately from autosomal variants. Quality control was conducted separately for males and females, using similar quality control parameters as above. Cases and pseudo-controls were built out of the trios. Phasing and imputation were then performed on males and females separately for each sample, followed by logistic regression with the same PCs, and meta-analysis combining samples (same parameters as the autosomal analyses). Results were generated for EAS Stage 1 samples and EUR-EAS combined samples (excluding BIX1, BIX2 and BIX3). EAS Stage 2, BIX1, BIX2 and BIX3 samples do not have chromosome X data and were therefore not analyzed.

### Genetic correlation and heritability

Schizophrenia heritabilities in the observed scale for samples of EUR and EAS ancestry were estimated from their summary statistics using the LDSC^21^. We converted the heritabilities in the observed scale to liability scale assuming the schizophrenia population prevalence at 1%. The LD scores were pre-computed from the 1000 Genomes Project Phase 3 reference panel in EUR and EAS respectively (https://github.com/bulik/ldsc). Only autosomal variants having MAF greater than 5% in their respective population were included in the analysis, and variants in the MHC region were not included due to the long range LD.

We computed the genetic correlations between schizophrenia and other traits within EUR and across EUR and EAS. EUR and EAS (Stage 1 only) summary statistics for autosomal variants from this study were used as schizophrenia genetic association inputs for their respective populations. Traits tested included schizophrenia^2^, bipolar^67^, major depression^68^, anorexia nervosa^69^, neuroticism^70^, autism spectrum disorder (PGC 2015 release), attention deficit hyperactivity disorder (with samples of non-European ancestry removed, available at http://www.med.unc.edu/pgc)^71^, education attainment^72^, general intelligence^73^, fluid intelligence score and prospective memory result (using individuals from UK Biobank), and subjective well being (SWB)^70^. Only variants having MAF greater than 5% were available and included.

Variants in the MHC region were excluded from the analysis. Genetic correlations within EUR were computed using LDSC with LD scores pre-computed on the 1000 Genomes Project Phase 3 reference panel (503 EUR subjects). Genetic correlations across EUR and EAS were computed using POPCORN^27^. POPCORN uses a Bayesian approach which assumes that genotypes are drawn separately from each population and effect sizes follow the infinitesimal model. The inflation of *z* scores could then be modelled and a weighted likelihood function which was maximized to find heritability and genetic correlation. Genetic correlations in POPCORN were computed in the “genetic effect” mode, which estimates the correlation based on the LD covariance scores and effect sizes from summary statistics.

### Partitioned heritability

Partitioned LDSC^32^ was conducted to look for heritability enrichment in diverse annotations using EAS (Stage 1) and EUR autosomal variants (summary statistics) respectively. LD scores for each annotation were computed using a combination of PLINK^64^ and LDSC^21^ using the 1000 Genomes Project EAS and EUR subjects respectively. We used baseline annotations^32^ and additional annotations including chromatin accessibility in brain dorso-lateral prefrontal cortex through the Assay for Transposase-Accessible Chromatin using sequencing peaks (ATAC Bryois)^33^, conserved regions located in “ATAC Bryois” (ATAC Bryois & Conserved LindbladToh)^33^, and introgressed regions from Neanderthal (Neanderthal Vernot)^74^. Variants can be included in multiple annotations. Multi-allelic variants were removed.

### Gene-set analysis

We performed gene and gene-set based tests using MAGMA^35^. Genome-wide summary statistics for autosomal variants from EAS, EUR and EAS+EUR meta-analyses were used in this analysis. Variant-to-gene annotation was performed using RefSeq NCBI37.3 with a window of 5 Kb upstream and 1.5 Kb downstream. LD was taken from 1000 Genomes Project EAS, EUR and EUR-EAS panels respectively. The gene-based *P*-values were computed using *F*-test and multivariate linear model, and competitive tests were used for gene-set analysis. Seventy gene-sets were selected and tested in this study (Supplementary Table 7) including those from the Molecular Signatures Database databases^75^, related to psychiatric diseases^36,76,77^ and from ‘gwaspipeline’ (http://github.com/freeseek/gwaspipeline/blob/master/makegenes.sh). Genesets were ranked for EUR, EAS and EAS+EUR analyses respectively. The top ranking genesets were compared across analyses to identify common schizophrenia pathways. Additionally, Wilcoxon sign rank tests was conducted to compare the ranking of gene-sets between the EUR and EAS datasets.

### Natural selection analysis

We used the CHB and CEU panels from the 1000 Genomes Project Phase 3 to investigate the natural selection signatures in schizophrenia-associated loci for EAS and EUR populations respectively. We used the following selection signatures, with their sensitivity to timeframes discussed in ref 3. *integrated Haplotype Score (iHS)*: iHS captures the haplotype homozygosity at a given variant. We calculated iHS using the R rehh package^78^. Genetic distance between variants was determined using HapMap phase II genetic map. Ancestral and derived alleles were obtained from the 1000 Genome project, which inferred the ancestral state using six primates on the EPO (Enredo-Pecan-Ortheus) pipeline. Only bi-allelic variants that have MAF ≥ 5% were included in the analysis. *Cross Population Extended Haplotype Homozygosity (XPEHH)^79^*: XPEHH detects variants under selection in one population but not the other. We used CEU as the reference panel when calculating XPEHH for CHB and vice versa. *Fixation index (Fst)*: Fst measures the population differentiation due to genetic structure. We estimated *Fst* using the Weir and Cockerham approach^80^, which is robust to sample size effects. *Absolute derived allele frequency difference (|ΔDAF|)*: |ΔDAF| measures population differentiation between CHB and CEU populations. *Composite of Multiple signals (CMS)^81-83^*: CMS combines iHS, XPEHH, *Fst* and |ΔDAF|. As a result, CMS potentially has better power to detect the selection signature. For each variant, 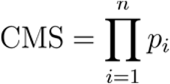, in which *P_i_* is the rank of the variant using method *i,* sorted by increasing *P*-values, divided by the total number of variants. *B statistic:* B statistic measures the background selection. We calculated the B statistic as in ref 84.

#### Trans-ethnicity fine-mapping

For a disease-associated genetic locus, fine-mapping defines a “credible set” of variants that contains the causal variant with certain probability (e.g., 99% or 95%). The Bayesian fine-mapping approaches^2,38,85,86^ have been widely used for studies of a single ancestry. Here, we extended a Bayesian fine-mapping approach^85^ (Defining credible sets, Methods) to studies of more than one ancestry.

Assume *D* represents the data including the genotype matrix *X* for all the *P* variants and disease status *Y* for *N* individuals, and *β* represents a collection of model parameters. We define the model, denoted by *A,* as the causal status for the *P* variants in locus: *A* ≡ {*a_j_*}, in which *a_j_* is the causal status for variant *j. a_j_* = 1 if the variant *j* is causal, and *a_j_* = 0 if it is not. We assume that there is one and only one genuine signal for each locus, and the causal variant is the same across all ancestries; therefore, one and only one of the *P* variants is causal: 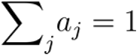. For convenience, we define *^A^_j_* as the model in which only variant *j* is causal, and as the model in which no variant is causal (null model). The probability of model *^A^_j_* (where variant *j* is the only causal variant in the locus) given the data *(D)* can be calculated using Bayes’s rule:

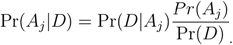

Wth the steepest descent approximation, the assumption of a flat prior on the model parameters (*β*), and the assumption of one causal variant per locus (equation 2 in ref 85), Pr(*A_j_*|*D*) can be approximated as:

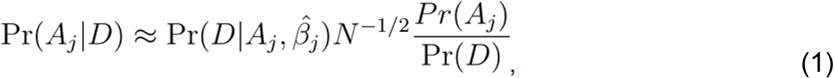

in which *N* is the sample size. We denote 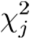 as the *x^2^* test statistic for variant *j*, which can be calculated from the *P*-value from the meta-analysis combining EAS and EUR samples. Using equation 3 in ref 85, we have

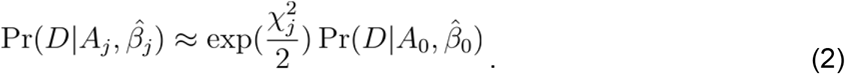

Pr(*A_j_*) is the prior probability that variant *j* is causal. We have shown that schizophrenia causal variants have consistent genetic effect across populations. Therefore we model the prior probability as a function of the heterogeneity measured in *I^2^*:

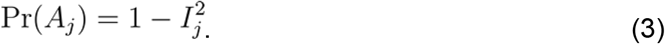

Using equations 2 and 3, Pr(*A_j_*|*D*) in equation 1 can be calculated as

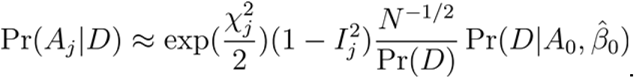

We only use Stage 1 samples in fine-mapping so the variants have the same sample size (assuming all variants have good imputation quality). Therefore, *N*^−1/2^,Pr(*D*) and 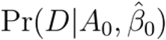 can be regarded as constants,

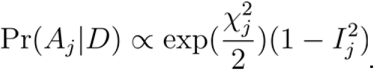

The normalized causal probability for variant *j* is then

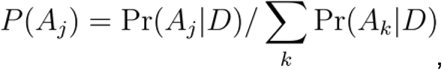

and the 95% credit set of variants is defined as the smallest set of variants, *S*, such that

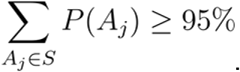

#### Polygenic risk score analysis

We constructed PRS using a pruning and thresholding approach in a study set of EAS individuals with training summary statistics from either EUR or EAS individuals. In the former case, we used summary statistics from all EUR individuals in this study; in the latter case, we used a leave-one-out meta-analysis approach across the 13 Stage 1 samples to build PRS.

For the EUR training data, we extracted EUR individuals (FIN, GBR, CEU, IBS, TSI) from 1000 Genomes Project^63^ Phase 3 as an LD reference panel to greedily clump variants. For the EAS LD reference panel, we created two panels: 1) an analogous EAS panel (CDX, CHB, CHS, JPT, KHV) from 1000 Genome Project^63^ Phase 3 (Fig. 4 and Extended Data Fig. c and d), and 2) an LD panel from best guess genotypes from each cohort in the study (Extended Data Fig. a,b,e,f). For both EAS and EUR prediction sets, we filtered to variants with a MAF greater than 1% in each respective populations, and removed indels and strand ambiguous variants.

We subset each list of variants to those in the summary statistics with an imputation INFO > 0.9. We then selected approximately independent loci at varying P-value thresholds or top-ranking *n* variants using an LD threshold of *R^2^* ≤ 0.1 in a window of 500 kilobase pairs in PLINK^64^ with the --clump flag. We treated the MHC with additional caution to minimize overfitting in this region, selecting only the most significant variant from the HLA region. To profile variants, we multiplied the log odds ratio for selected variants by genotypes and summed these values across the genome in PLINK^64^ using the --score flag for each of the 13 EAS Stage 1 samples. We assessed case/control variance explained by computing Nagelkerke’s and a liability-scale pseudo-*R*^2^ as in Lee *et al.^87^* by comparing a full model with the PRS and 10 principal components with a model excluding the PRS.

#### Data availability

Summary statistics from this study can be downloaded from http://personal.broadinstitute.org/hhuang/PGC_SCZ_EAS/. Raw genotype data that support the findings of this study are available from the Psychiatric Genomics Consortium but restrictions apply to the availability of these data, which were used under licence for the current study, and so are not publicly available. Data are, however, available from the corresponding authors upon reasonable request and with the permission of the Psychiatric Genomics Consortium.

#### Code availability

Computer code used to perform QC, PCA, imputation, association test and meta-analysis can be downloaded from http://github.com/Nealelab/ricopili/wiki. Code for other analyses is available upon request.

---

**Extended Data Figure 1.**
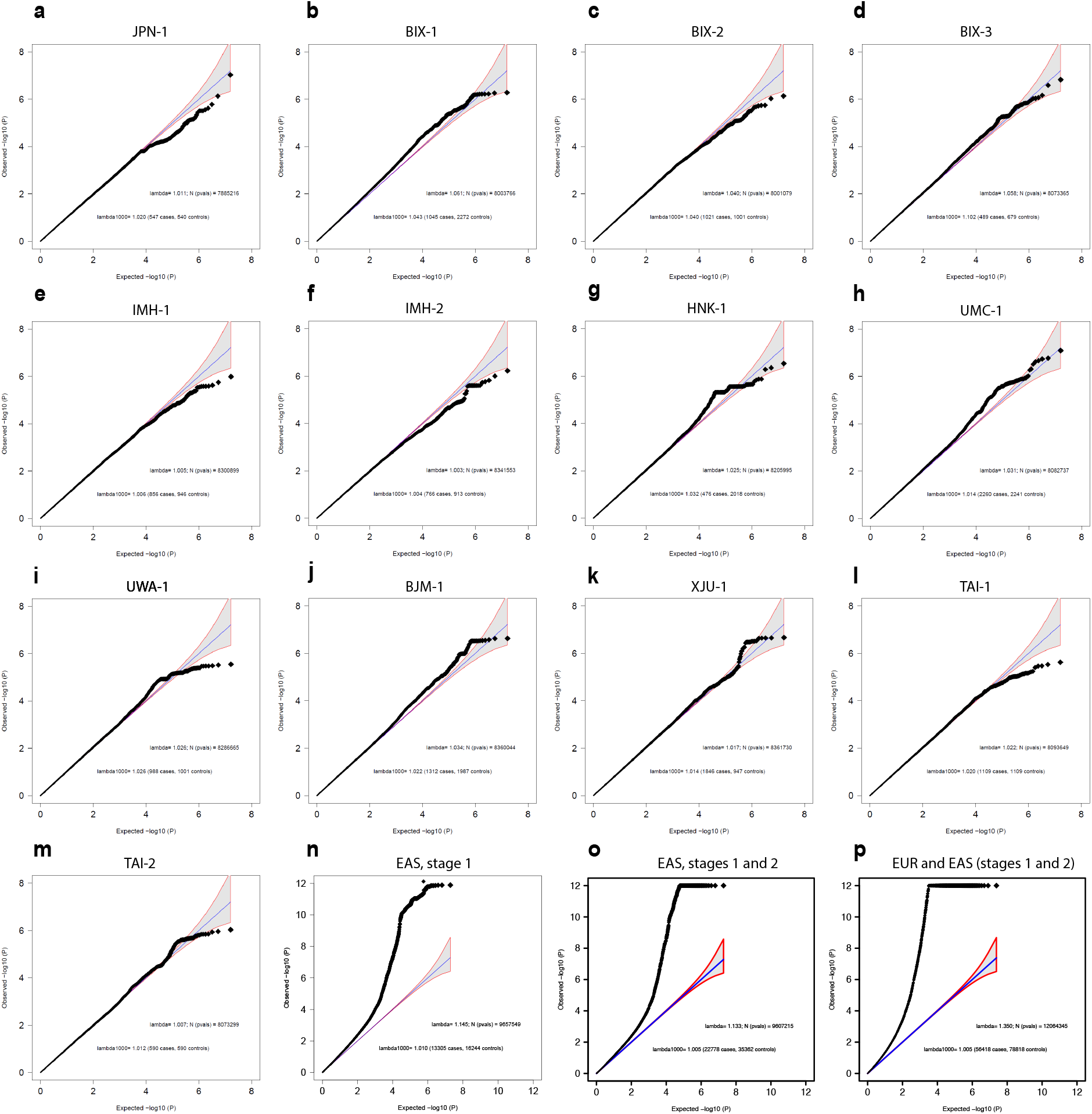
Quantile-quantile (QQ) plots. QQ plots for each EAS stage-1 samples (**a-m**) and meta-analyses including all EAS Stage 1 samples (**n**), Stages 1 and 2 samples (**o**) and all EUR and EAS (Stages 1 and 2) samples (**p**). Blue line indicates the expected null distribution, and the shaded area indicates the 95% confidence interval of the null distribution. Legend: “lambda”=genomic inflation factor; “lamda1000”=genomic inflation factor for an equivalent study of 1000 cases and 1000 controls; and “N(pvals)”=number of variants used in the plot. Autosomal variants that have minor allele frequency ≥ 1% and INFO ≥ 0.6 from imputation were included. Observed P-values were capped at 10^-12^ for visualization purpose.

**Extended Data Figure 2.**
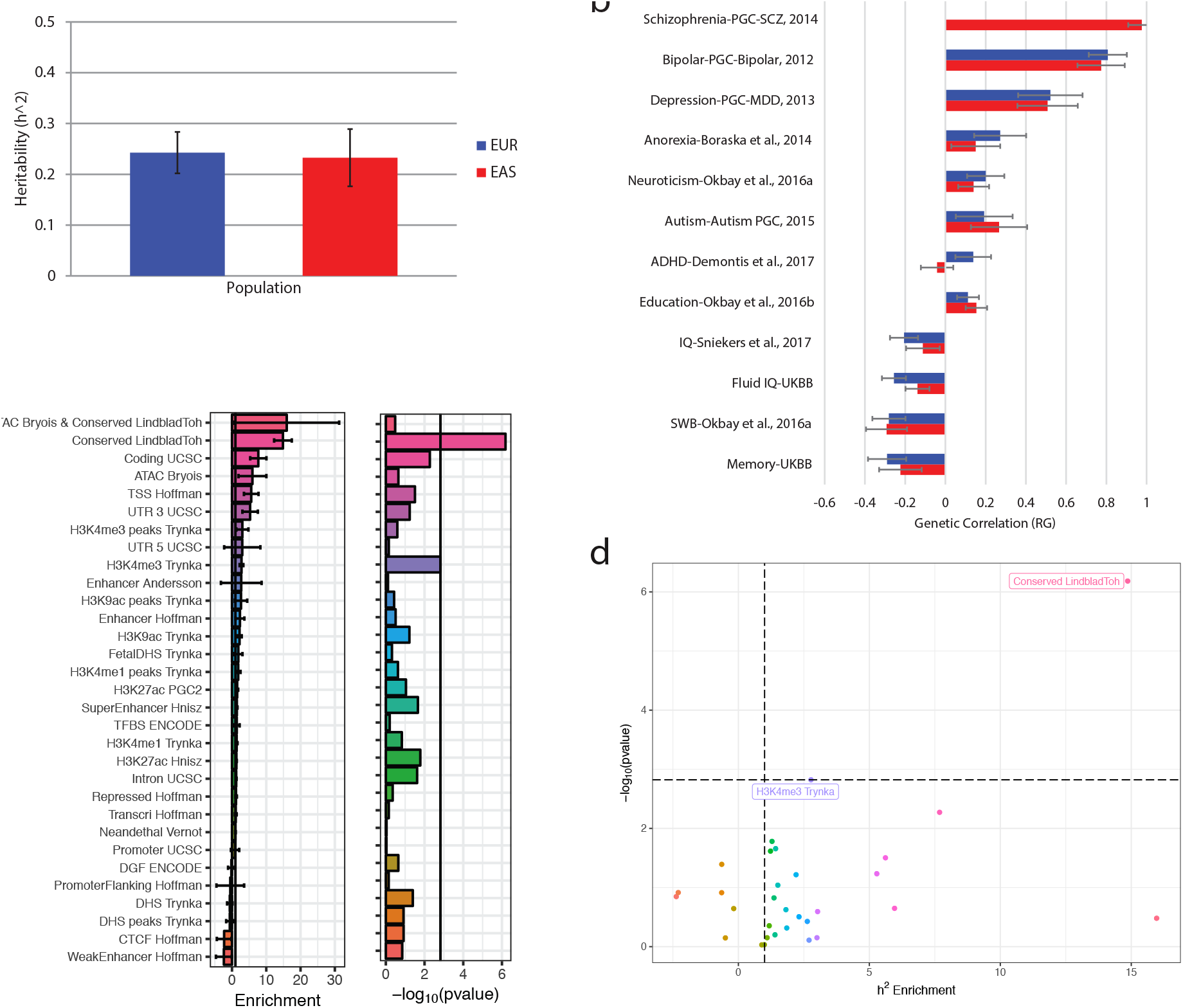
Heritability and genetic correlation. **a,** Heritability (h^2^) for the EAS and EUR samples. **b,** Genetic correlation between schizophrenia and other traits within EUR (blue) and across EAS and EUR (red). Error bars indicate the 95% confidence interval. **c,** Enrichment and its corresponding significance for heritability partitioned based on various annotations. **d,** Scatterplot showing the enrichment versus the significance for heritability partitioned based on various annotations. More details are available in Methods.

**Extended Data Figure 3.**
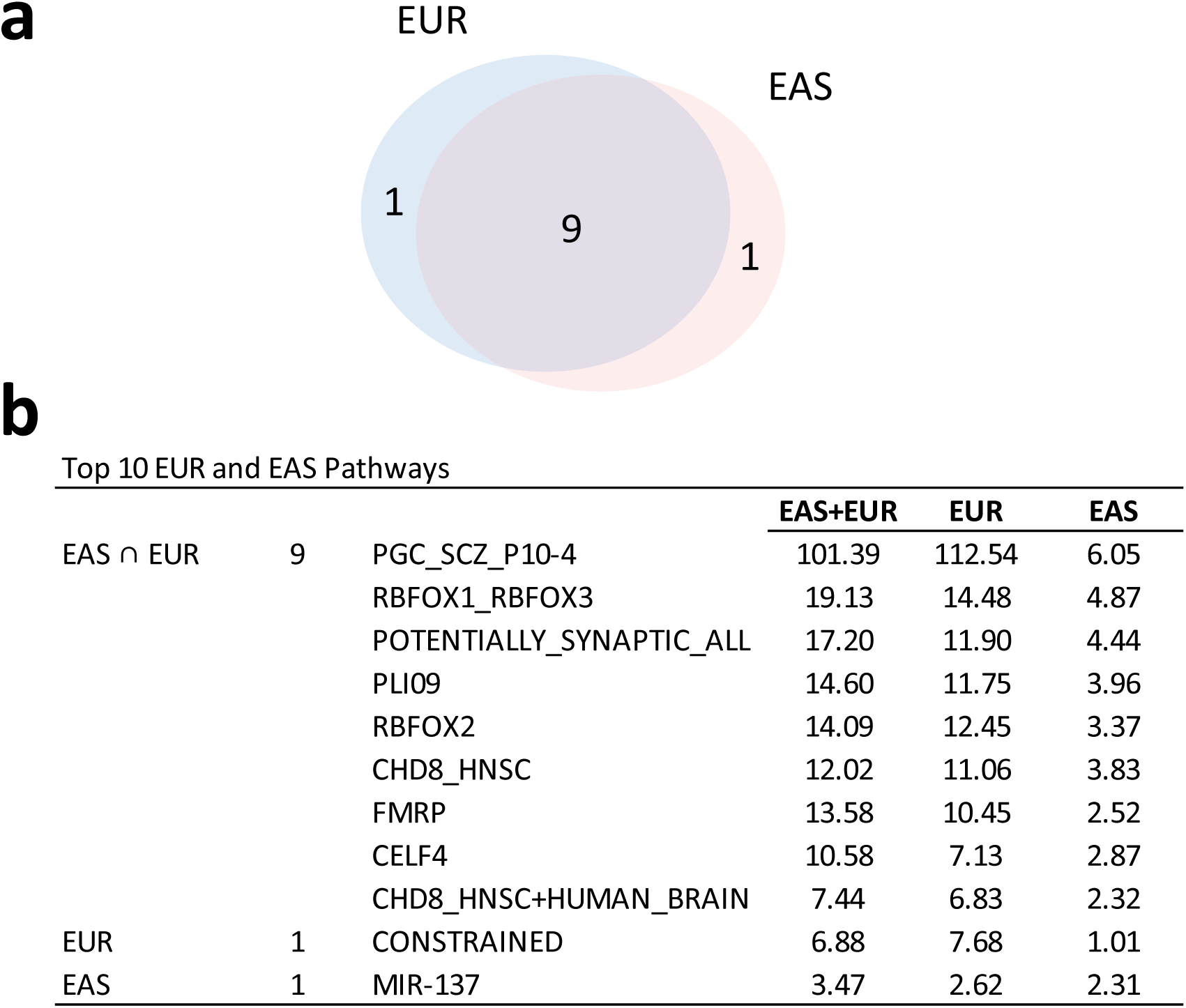
Gene-sets implicated by schizophrenia genetic associations. **a,** Overlap of implicated gene-sets across EUR and EAS samples. **b,** List of the top 10 gene-sets implicated in the EAS and EUR samples and their *P*-values in -log_10_ scale. Descriptions of the gene-sets are available in Supplementary Table 8.

**Extended Data Figure 4.**
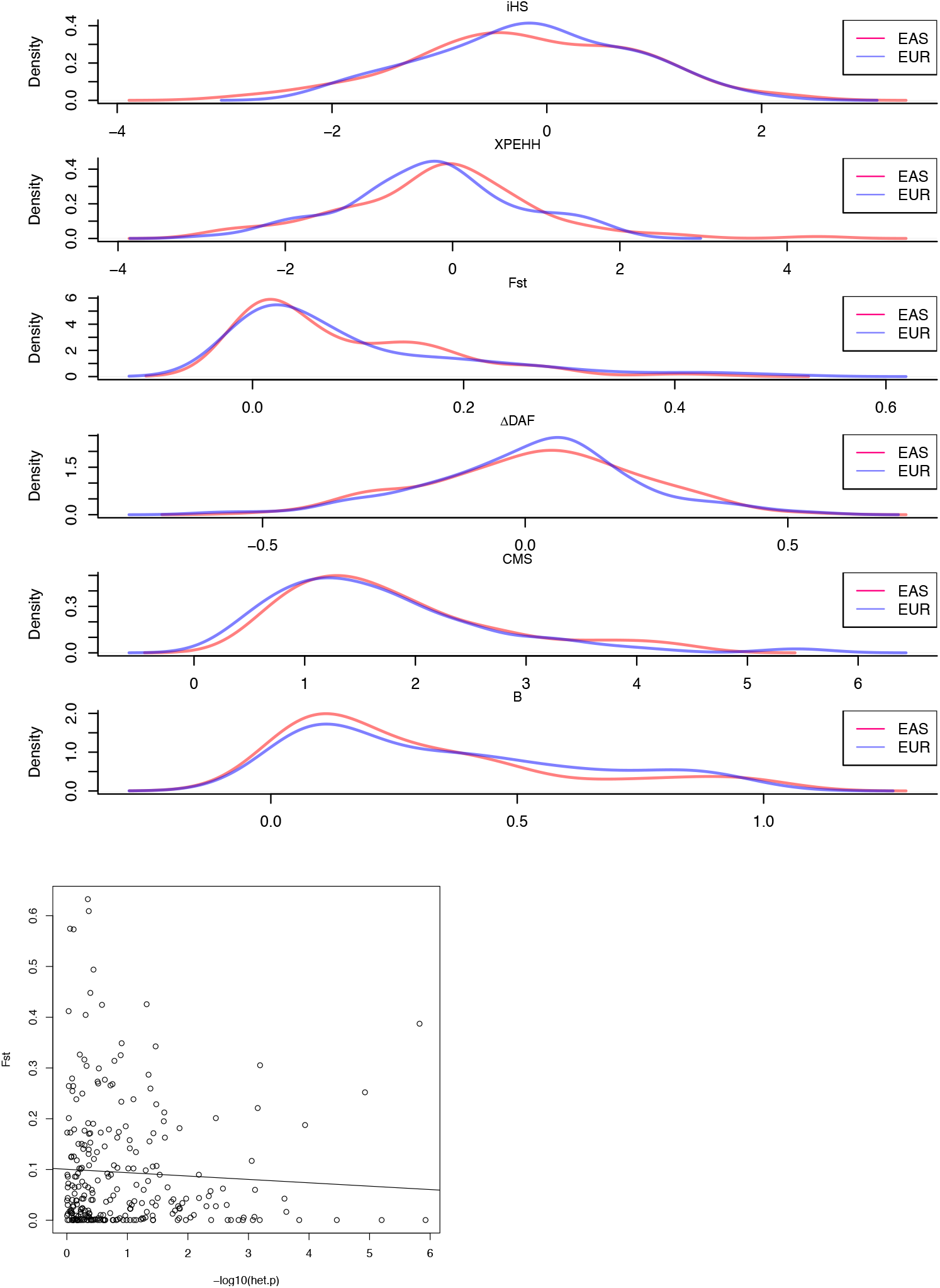
Natural selection signals in EAS and EUR. **a,** Distributions of natural selection signals in the top 100 schizophrenia associations in EAS (red) and EUR (blue). **b,** Scatterplot of *Fst* versus the heterogeneity of effect size for schizophrenia associations. More details are available in Methods.

**Extended Data Figure 5.**
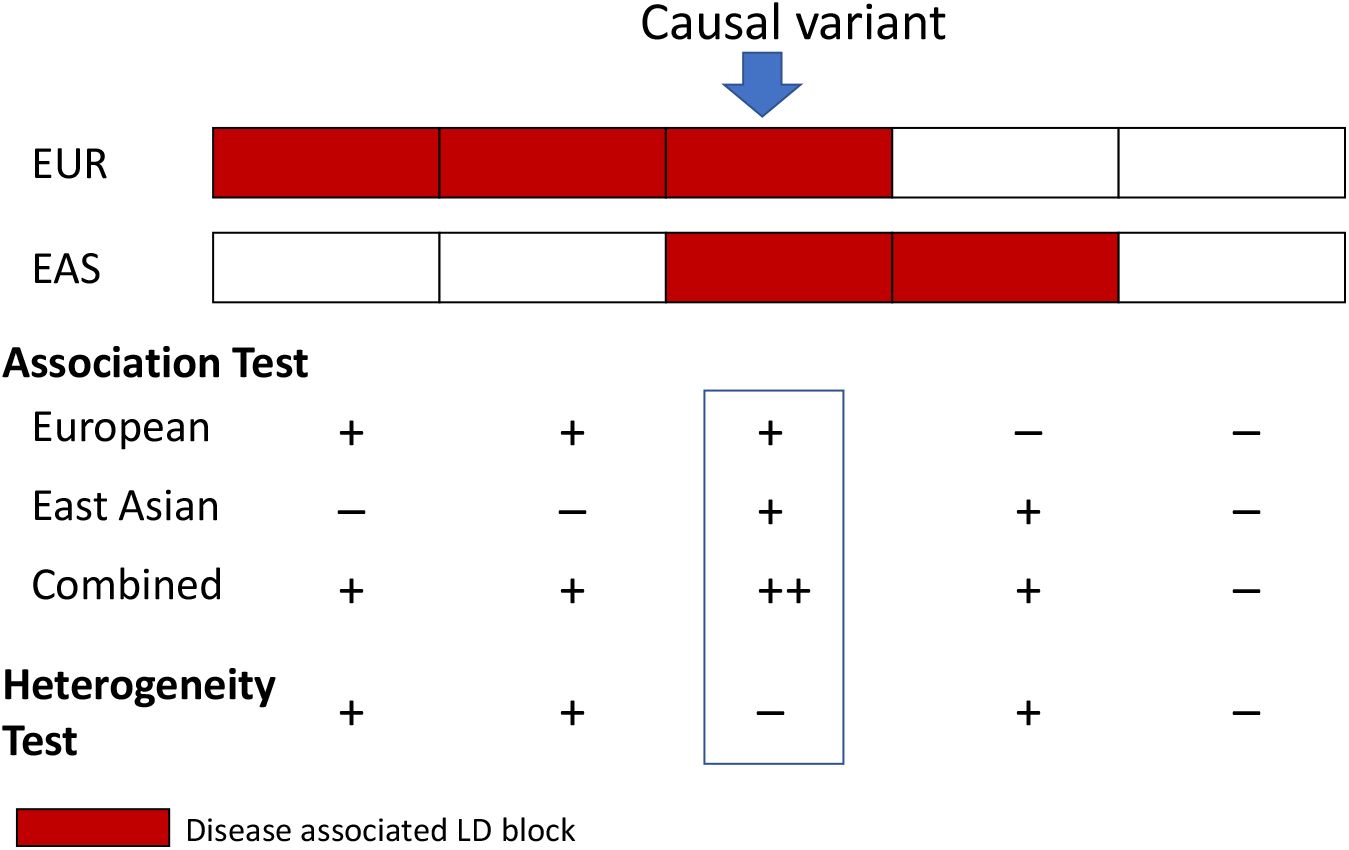
Trans-ethnicity fine-mapping. Illustration of the fine-mapping method.

**Extended Data Figure 6.**
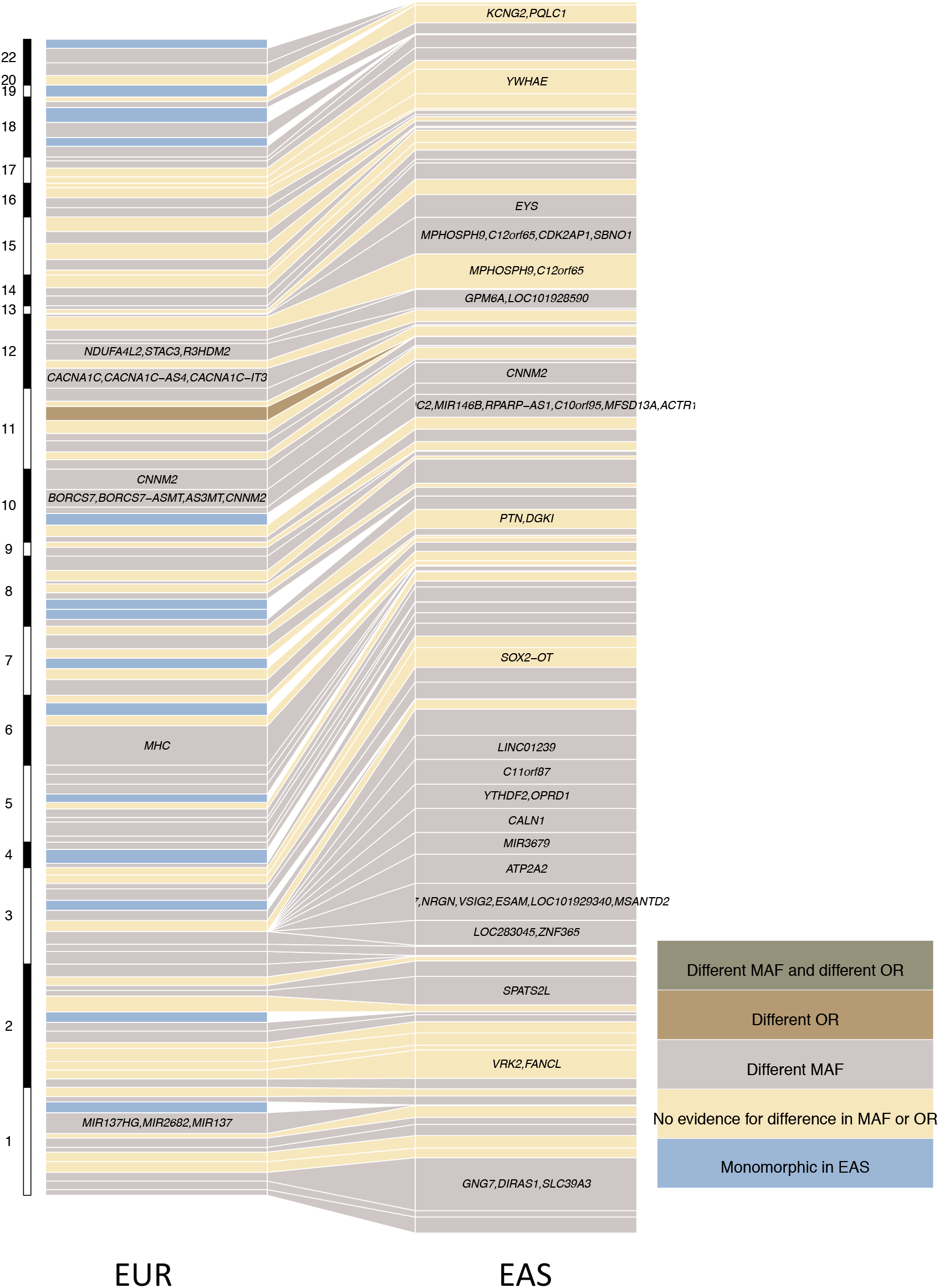
Variance explained for schizophrenia associations across EUR and EAS samples. Genome-wide significant associations that have variance explained greater than 0.05% in either EAS or EUR samples were plotted. One locus can host multiple independent associations. Different MAF is defined as *Fst*>0.01, and different OR is defined as heterogeneity test *P*-value < 0.05 after bonferroni correction. Nearest genes to the associations were used as labels for associations when the text space is available, with the exception that the MHC locus was labeled as “MHC”.

**Extended Data Figure 7.**
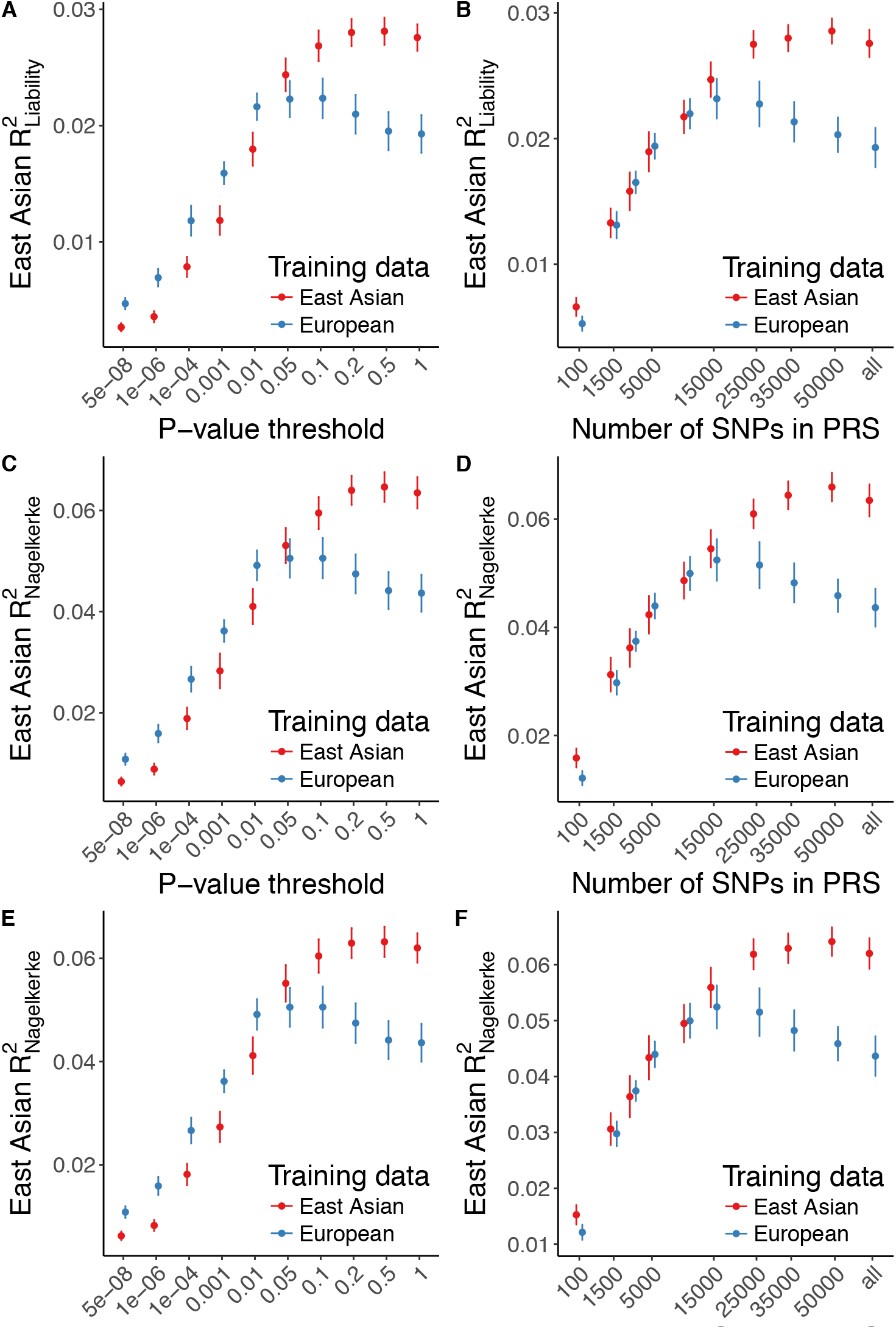
Genetic risk prediction accuracy in EAS from EAS or EUR training data. As in Fig. 4, PRS shows case/control variance explained with EUR and EAS samples using a leave-one-out meta-analysis approach for the EAS samples. Error bars indicate the 95% confidence intervals. **a,b)** Liability-scale variance explained when LD panel for clumping is from EUR 1000 Genomes Phase 3 samples and best-guess genotypes are from each EAS cohort. **c,d)** Nagelkerke’s *R*^2^ for EAS prediction accuracy when LD panel for clumping is from EUR and EAS 1000 Genomes Phase 3 samples. **E-F)** Nagelkerke’s *R*^2^ for EAS prediction accuracy when LD panel for clumping is from EUR 1000 Genomes Phase 3 samples and best-guess genotypes are from each EAS cohort.

**Extended Data Figure 8.**
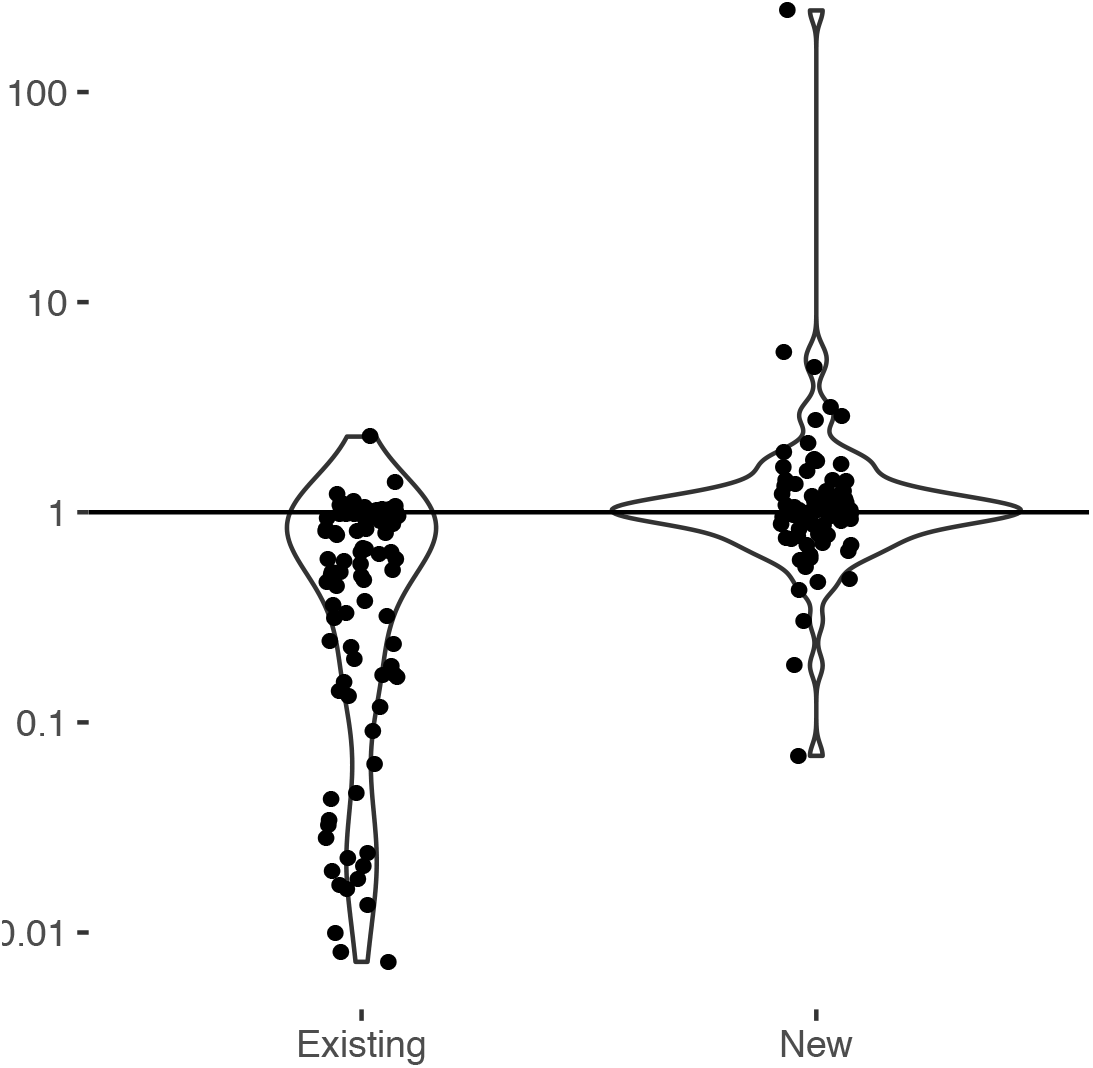
Ratio of the heterozygote rate in EAS to that in EUR for existing and new loci. Het(EAS) and Het(EUR), calculated as 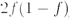, are the heterozygote rates for a variant in EAS and EUR respectively, in which 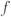 is the variant allele frequency in EAS or EUR. Power to identify genetic associations increases with the expected non-centrality parameter for the association, which is proportional to the heterozygote rate. Therefore we use the ratio of the heterozygote rate in EAS to that in EUR as a measure of the relative power to identify genetic association of the same effect size in the two populations. A ratio greater than 1 means EAS samples have more power to identify the association and vice versa. Existing loci are those that are genome-wide significant in the previous study of European ancestry^2^, and new loci are those that are genome-wide significant just in this study combining EAS and EUR samples.

**Extended Data Figure 9.**
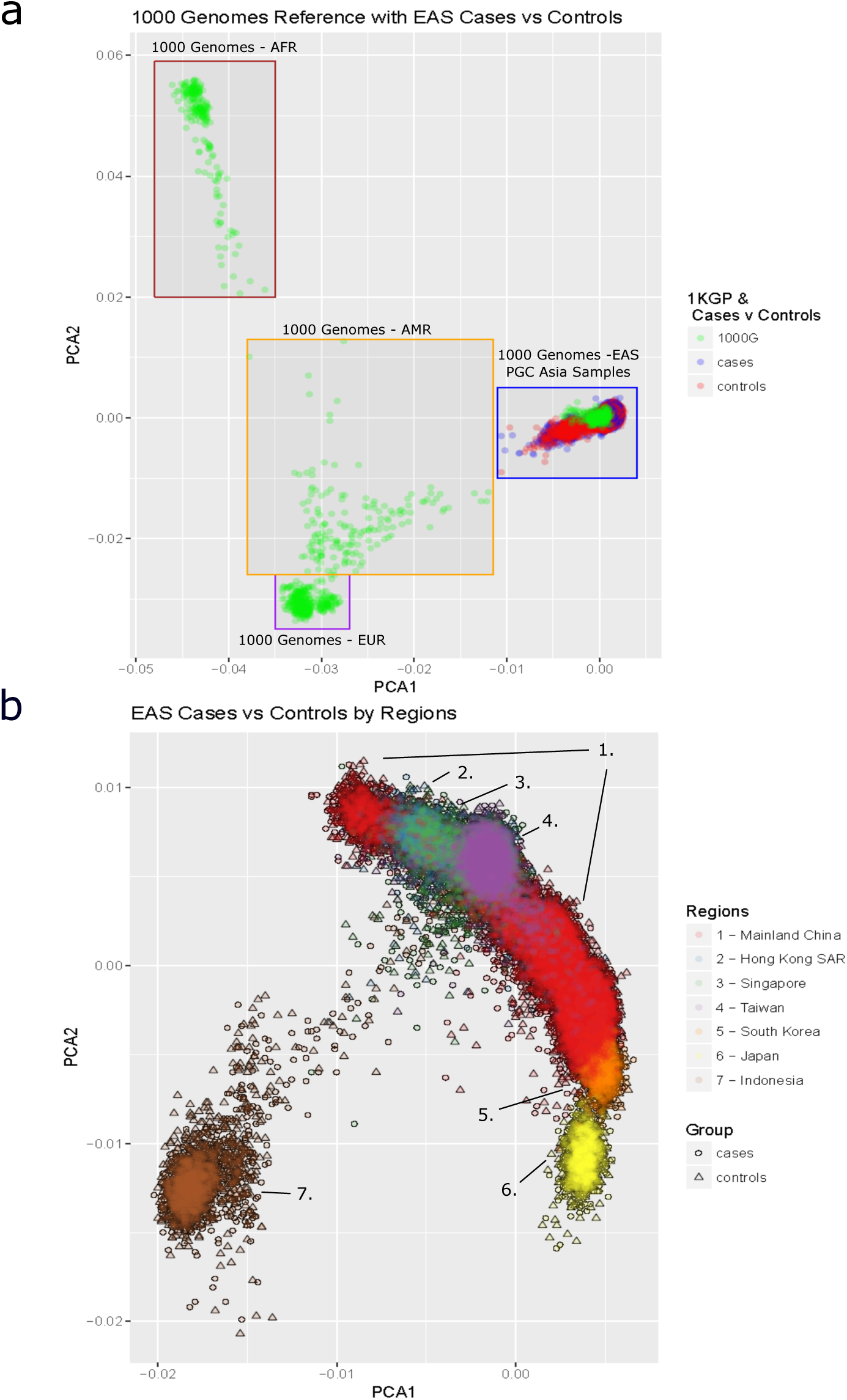
Extended Data Figure 9Principal component analysis of EAS samples. **a,** EAS samples mapped to the global principal components created using 1000 Genomes Project Phase 1 samples. **b,** EAS cases and controls mapped respectively to principal components created using all EAS samples in this study.

**Extended Data Table 1.**
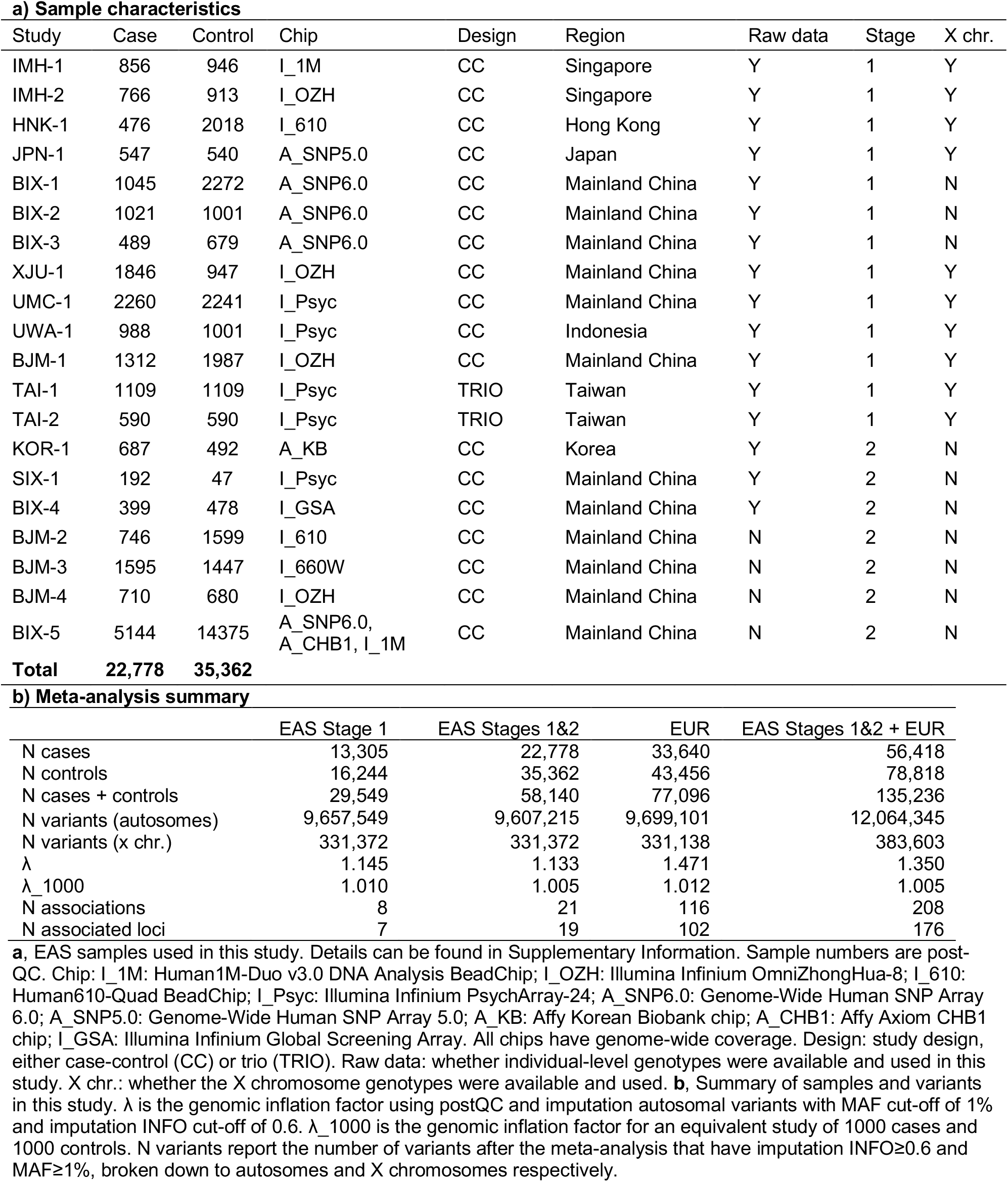
Overview of samples and variants

a) Sample characteristics
| Study | Case | Control | Chip | Design | Region | Raw data | Stage | X chr. |
| --- | --- | --- | --- | --- | --- | --- | --- | --- |
| IMH-1 | 856 | 946 | I_1M | CC | Singapore | Y | 1 | Y |
| IMH-2 | 766 | 913 | I_OZH | CC | Singapore | Y | 1 | Y |
| HNK-1 | 476 | 2018 | I_610 | CC | Hong Kong | Y | 1 | Y |
| JPN-1 | 547 | 540 | A_SNP5.0 | CC | Japan | Y | 1 | Y |
| BIX-1 | 1045 | 2272 | A_SNP6.0 | CC | Mainland China | Y | 1 | N |
| BIX-2 | 1021 | 1001 | A_SNP6.0 | CC | Mainland China | Y | 1 | N |
| BIX-3 | 489 | 679 | A_SNP6.0 | CC | Mainland China | Y | 1 | N |
| XJU-1 | 1846 | 947 | I_OZH | CC | Mainland China | Y | 1 | Y |
| UMC-1 | 2260 | 2241 | I_Psyc | CC | Mainland China | Y | 1 | Y |
| UWA-1 | 988 | 1001 | I_Psyc | CC | Indonesia | Y | 1 | Y |
| BJM-1 | 1312 | 1987 | I_OZH | CC | Mainland China | Y | 1 | Y |
| TAI-1 | 1109 | 1109 | I_Psyc | TRIO | Taiwan | Y | 1 | Y |
| TAI-2 | 590 | 590 | I_Psyc | TRIO | Taiwan | Y | 1 | Y |
| KOR-1 | 687 | 492 | A_KB | CC | Korea | Y | 2 | N |
| SIX-1 | 192 | 47 | I_Psyc | CC | Mainland China | Y | 2 | N |
| BIX-4 | 399 | 478 | I_GSA | CC | Mainland China | Y | 2 | N |
| BJM-2 | 746 | 1599 | I_610 | CC | Mainland China | N | 2 | N |
| BJM-3 | 1595 | 1447 | I_660W | CC | Mainland China | N | 2 | N |
| BJM-4 | 710 | 680 | I_OZH | CC | Mainland China | N | 2 | N |
| BIX-5 | 5144 | 14375 | A_SNP6.0,<br>A_CHB1, I_1M | CC | Mainland China | N | 2 | N |
| **Total** | **22,778** | **35,362** |  |  |  |  |  |  |

b) Meta-analysis summary
|  | EAS Stage 1 | EAS Stages 1&2 | EUR | EAS Stages 1&2 + EUR |
| --- | --- | --- | --- | --- |
| N cases | 13,305 | 22,778 | 33,640 | 56,418 |
| N controls | 16,244 | 35,362 | 43,456 | 78,818 |
| N cases + controls | 29,549 | 58,140 | 77,096 | 135,236 |
| N variants (autosomes) | 9,657,549 | 9,607,215 | 9,699,101 | 12,064,345 |
| N variants (x chr.) | 331,372 | 331,372 | 331,138 | 383,603 |
| $\lambda$ | 1.145 | 1.133 | 1.471 | 1.350 |
| $\lambda_{1000}$ | 1.010 | 1.005 | 1.012 | 1.005 |
| N associations | 8 | 21 | 116 | 208 |
| N associated loci | 7 | 19 | 102 | 176 |

**a**, EAS samples used in this study. Details can be found in Supplementary Information. Sample numbers are post-QC. Chip: I\_1M: Human1M-Duo v3.0 DNA Analysis BeadChip; I\_OZH: Illumina Infinium OmniZhongHua-8; I\_610: Human610-Quad BeadChip; I\_Psyc: Illumina Infinium PsychArray-24; A\_SNP6.0: Genome-Wide Human SNP Array 6.0; A\_SNP5.0: Genome-Wide Human SNP Array 5.0; A\_KB: Affy Korean Biobank chip; A\_CHB1: Affy Axiom CHB1 chip; I\_GSA: Illumina Infinium Global Screening Array. All chips have genome-wide coverage. Design: study design, either case-control (CC) or trio (TRIO). Raw data: whether individual-level genotypes were available and used in this study. X chr.: whether the X chromosome genotypes were available and used. **b**, Summary of samples and variants in this study. $\lambda$ is the genomic inflation factor using postQC and imputation autosomal variants with MAF cut-off of 1% and imputation INFO cut-off of 0.6. $\lambda_{1000}$ is the genomic inflation factor for an equivalent study of 1000 cases and 1000 controls. N variants report the number of variants after the meta-analysis that have imputation INFO $\geq 0.6$ and MAF $\geq 1\%$ , broken down to autosomes and X chromosomes respectively.

## References

1. Stilo, S. A. & Murray, R. M. The epidemiology of schizophrenia: replacing dogma with knowledge. Dialogues Clin. Neurosci. 12, 305–315 (2010).

2. Schizophrenia Working Group of the Psychiatric Genomics Consortium. Biological insights from 108 schizophrenia-associated genetic loci. Nature 511, 421–427 (2014).

3. Pardiñas, A. F. et al. Common schizophrenia alleles are enriched in mutation-intolerant genes and in regions under strong background selection. Nat. Genet. (2018). doi:10.1038/s41588-018-0059-2

4. Visscher, P. M. et al. 10 Years of GWAS Discovery: Biology, Function, and Translation. Am. J. Hum. Genet. 101, 5–22 (2017).

5. Li, Y. R. & Keating, B. J. Trans-ethnic genome-wide association studies: advantages and challenges of mapping in diverse populations. Genome Med. 6, 91 (2014).

6. Popejoy, A. B. & Fullerton, S. M. Genomics is failing on diversity. Nature 538, 161–164 (2016).

7. Yu, H. et al. Common variants on 2p16.1, 6p22.1 and 10q24.32 are associated with schizophrenia in Han Chinese population. Mol. Psychiatry 22, 954–960 (2017).

8. Li, Z. et al. Genome-wide association analysis identifies 30 new susceptibility loci for schizophrenia. Nat. Genet. (2017). doi:10.1038/ng.3973

9. Moltke, I. et al. A common Greenlandic TBC1D4 variant confers muscle insulin resistance and type 2 diabetes. Nature 512, 190–193 (2014).

10. Tang, C. S. et al. Exome-wide association analysis reveals novel coding sequence variants associated with lipid traits in Chinese. Nat. Commun. 6, 10206 (2015).

11. Clarke, T.-K. et al. Genome-wide association study of alcohol consumption and genetic overlap with other health-related traits in UK Biobank (N=112 117). Mol. Psychiatry 22, 1376–1384 (2017).

12. CONVERGE consortium. Sparse whole-genome sequencing identifies two loci for major depressive disorder. Nature 523, 588–591 (2015).

13. Wray, N. R. et al. Genome-wide association analyses identify 44 risk variants and refine the genetic architecture of major depression. Nat. Genet. 50, 668–681 (2018).

14. World Health Organization. Report of the International Pilot Study of Schizophrenia. (1973).

15. Carpenter, W. T., Jr, Strauss, J. S. & Bartko, J. J. Flexible system for the diagnosis of schizophrenia: report from the WHO International Pilot Study of Schizophrenia. Science 182, 1275–1278 (1973).

16. Jablensky, A. et al. Schizophrenia: manifestations, incidence and course in different cultures. A World Health Organization ten-country study. Psychol. Med. Monogr. Suppl. 20, 1–97 (1992).

17. McGrath, J. J. Variations in the incidence of schizophrenia: data versus dogma. Schizophr. Bull. 32, 195–197 (2006).

18. Haro, J. M. et al. Cross-national clinical and functional remission rates: Worldwide Schizophrenia Outpatient Health Outcomes (W-SOHO) study. Br. J. Psychiatry 199, 194-201 (2011).

19. Gureje, O. & Cohen, A. Differential outcome of schizophrenia: where we are and where we would like to be. Br. J. Psychiatry 199, 173–175 (2011).

20. Martin, A. R. et al. Human Demographic History Impacts Genetic Risk Prediction across Diverse Populations. Am. J. Hum. Genet. 100, 635–649 (2017).

21. Bulik-Sullivan, B. et al. LD Score Regression Distinguishes Confounding from Polygenicity in Genome-Wide Association Studies. Nat. Genet. 47, 291–295 (2015).

22. Geisler, S., Schöpf, C. L. & Obermair, G. J. Emerging evidence for specific neuronal functions of auxiliary calcium channel a_2_δ subunits. Gen. Physiol. Biophys. 34, 105–118 (2015).

23. Heyes, S. et al. Genetic disruption of voltage-gated calcium channels in psychiatric and neurological disorders. Prog. Neurobiol. 134, 36–54 (2015).

24. Cross-Disorder Group of the Psychiatric Genomics Consortium. Identification of risk loci with shared effects on five major psychiatric disorders: a genome-wide analysis. Lancet 381, 1371–1379 (2013).

25. Gulsuner, S. et al. Spatial and temporal mapping of de novo mutations in schizophrenia to a fetal prefrontal cortical network. Cell 154, 518–529 (2013).

26. Liu, J. Z. et al. Association analyses identify 38 susceptibility loci for inflammatory bowel disease and highlight shared genetic risk across populations. Nat. Genet. 47, 979–986 (2015).

27. Brown, B. C., Asian Genetic Epidemiology Network Type 2 Diabetes Consortium, Ye, C. J., Price, A. L. & Zaitlen, N. Transethnic Genetic-Correlation Estimates from Summary Statistics. Am. J. Hum. Genet. 99, 76–88 (2016).

28. Okbay, A. et al. Genome-wide association study identifies 74 loci associated with educational attainment. Nature 533, 539–542 (2016).

29. Davies, G. et al. Genome-wide association study of cognitive functions and educational attainment in UK Biobank (N= 112 151). Mol. Psychiatry (2016).

30. Cross-Disorder Group of the Psychiatric Genomics Consortium et al. Genetic relationship between five psychiatric disorders estimated from genome-wide SNPs. Nat. Genet. 45, 984–994 (2013).

31. Anttila, V. et al. Analysis of shared heritability in common disorders of the brain. bioRxiv 048991 (2017). doi:10.1101/048991

32. Finucane, H. K. et al. Partitioning heritability by functional annotation using genome-wide association summary statistics. Nat. Genet. 47, 1228–1235 (2015).

33. Bryois, J. et al. Evaluation Of Chromatin Accessibility In Prefrontal Cortex Of Schizophrenia Cases And Controls. bioRxiv 141986 (2017). doi:10.1101/141986

34. Lindblad-Toh, K. et al. A high-resolution map of human evolutionary constraint using 29 mammals. Nature 478, 476–482 (2011).

35. de Leeuw, C. A., Mooij, J. M., Heskes, T. & Posthuma, D. MAGMA: generalized gene-set analysis of GWAS data. PLoS Comput. Biol. 11, e1004219 (2015).

36. Genovese, G. et al. Increased burden of ultra-rare protein-altering variants among 4,877 individuals with schizophrenia. Nat. Neurosci. 19, 1433–1441 (2016).

37. Michael Borenstein, Larry V. Hedges, Julian P. T. Higgins, Hannah R. Rothstein. Introduction to Meta-Analysis. (John Wiley & Sons, Ltd, 2009).

38. Huang, H. et al. Fine-mapping inflammatory bowel disease loci to single-variant resolution. Nature 547, 173–178 (2017).

39. Farh, K. K.-H. et al. Genetic and epigenetic fine mapping of causal autoimmune disease variants. Nature 518, 337–343 (2015).

40. Swaminathan, B. et al. Fine mapping and functional analysis of the multiple sclerosis risk gene CD6. PLoS One 8, e62376 (2013).

41. Gaulton, K. J. et al. Genetic fine mapping and genomic annotation defines causal mechanisms at type 2 diabetes susceptibility loci. Nat. Genet. 47, 1415–1425 (2015).

42. Kichaev, G. & Pasaniuc, B. Leveraging Functional-Annotation Data in Trans-ethnic Fine-Mapping Studies. Am. J. Hum. Genet. 97, 260–271 (2015).

43. Lin, W. et al. Hepatic metal ion transporter ZIP8 regulates manganese homeostasis and manganese-dependent enzyme activity. J. Clin. Invest. 127, 2407–2417 (2017).

44. Park, J. H. et al. SLC39A8 Deficiency: A Disorder of Manganese Transport and Glycosylation. Am. J. Hum. Genet. 97, 894–903 (2015).

45. Li, D. et al. A Pleiotropic Missense Variant in SLC39A8 Is Associated With Crohn’s Disease and Human Gut Microbiome Composition. Gastroenterology 151, 724–732 (2016).

46. International Consortium for Blood Pressure Genome-Wide Association Studies et al. Genetic variants in novel pathways influence blood pressure and cardiovascular disease risk. Nature 478, 103–109 (2011).

47. Savage, J. E. et al. GWAS meta-analysis (N=279,930) identifies new genes and functional links to intelligence. bioRxiv 184853 (2017). doi:10.1101/184853

48. Zhang, S. et al. Base-Specific Mutational Intolerance Near Splice-Sites Clarifies Role Of Non-Essential Splice Nucleotides. bioRxiv 129312 (2017). doi:10.1101/129312

49. Pawitan, Y., Seng, K. C. & Magnusson, P. K. E. How many genetic variants remain to be discovered? PLoS One 4, e7969 (2009).

50. Márquez-Luna, C., Loh, P.-R., South Asian Type 2 Diabetes (SAT2D) Consortium, SIGMA Type 2 Diabetes Consortium & Price, A. L. Multiethnic polygenic risk scores improve risk prediction in diverse populations. Genet. Epidemiol. 41, 811–823 (2017).

51. Chen, C.-Y., Han, J., Hunter, D. J., Kraft, P. & Price, A. L. Explicit Modeling of Ancestry Improves Polygenic Risk Scores and BLUP Prediction. Genet. Epidemiol. 39, 427–438 (2015).

52. Salinas, Y. D., Wang, L. & DeWan, A. T. Multiethnic genome-wide association study identifies ethnic-specific associations with body mass index in Hispanics and African Americans. BMC Genet. 17, 78 (2016).

53. International Schizophrenia Consortium et al. Common polygenic variation contributes to risk of schizophrenia and bipolar disorder. Nature 460, 748–752 (2009).

54. Sekar, A. et al. Schizophrenia risk from complex variation of complement component 4. Nature 530, 177–183 (2016).

55. Yue, W.-H. et al. Genome-wide association study identifies a susceptibility locus for schizophrenia in Han Chinese at 11p11.2. Nat. Genet. 43, 1228–1231 (2011).

56. Shi, Y. et al. Common variants on 8p12 and 1q24.2 confer risk of schizophrenia. Nat. Genet. 43, 1224–1227 (2011).

57. Chen, J. Y. et al. Effects of Complement C4 Gene Copy Number Variations, Size Dichotomy, and C4A Deficiency on Genetic Risk and Clinical Presentation of Systemic Lupus Erythematosus in East Asian Populations. Arthritis Rheumatol 68, 1442–1453 (2016).

58. Hong, G. H. et al. Association of complement C4 and HLA-DR alleles with systemic lupus erythematosus in Koreans. J. Rheumatol. 21, 442–447 (1994).

59. Mahajan, A. et al. Fine-mapping of an expanded set of type 2 diabetes loci to single-variant resolution using high-density imputation and islet-specific epigenome maps. bioRxiv 245506 (2018). doi:10.1101/245506

## References

60. Sand, P. G. A lesson not learned: allele misassignment. Behav. Brain Funct. 3, 65 (2007).

61. O’Connell, J. et al. A general approach for haplotype phasing across the full spectrum of relatedness. PLoS Genet. 10, e1004234 (2014).

62. Howie, B., Fuchsberger, C., Stephens, M., Marchini, J. & Abecasis, G. R. Fast and accurate genotype imputation in genome-wide association studies through pre-phasing. Nat. Genet. 44, 955–959 (2012).

63. 1000 Genomes Project Consortium et al. A global reference for human genetic variation. Nature 526, 68–74 (2015).

64. Purcell, S. et al. PLINK: a tool set for whole-genome association and population-based linkage analyses. Am. J. Hum. Genet. 81, 559–575 (2007).

65. Price, A. L. et al. Principal components analysis corrects for stratification in genome-wide association studies. Nat. Genet. 38, 904–909 (2006).

66. Willer, C. J., Li, Y. & Abecasis, G. R. METAL: fast and efficient meta-analysis of genomewide association scans. Bioinformatics 26, 2190–2191 (2010).

67. Psychiatric GWAS Consortium Bipolar Disorder Working Group. Large-scale genome-wide association analysis of bipolar disorder identifies a new susceptibility locus near ODZ4. Nat. Genet. 43, 977–983 (2011).

68. Major Depressive Disorder Working Group of the Psychiatric GWAS Consortium et al. A mega-analysis of genome-wide association studies for major depressive disorder. Mol. Psychiatry 18, 497–511 (2013).

69. Boraska, V. et al. A genome-wide association study of anorexia nervosa. Mol. Psychiatry 19, 1085–1094 (2014).

70. Okbay, A. et al. Genetic variants associated with subjective well-being, depressive symptoms, and neuroticism identified through genome-wide analyses. Nat. Genet. 48, 624-633 (2016).

71. Demontis, D. et al. Discovery Of The First Genome-Wide Significant Risk Loci For ADHD. bioRxiv 145581 (2017). doi:10.1101/145581

72. Okbay, A. et al. Genome-wide association study identifies 74 loci associated with educational attainment. Nature 533, 539–542 (2016).

73. Sniekers, S. et al. Genome-wide association meta-analysis of 78,308 individuals identifies new loci and genes influencing human intelligence. Nat. Genet. 49, 1107–1112 (2017).

74. Juric, I., Aeschbacher, S. & Coop, G. The Strength of Selection against Neanderthal Introgression. PLoS Genet. 12, e1006340 (2016).

75. Liberzon, A. et al. Molecular signatures database (MSigDB) 3.0. Bioinformatics 27, 1739-1740 (2011).

76. Turner, T. N. et al. denovo-db: a compendium of human de novo variants. Nucleic Acids Res. 45, D804–D811 (2017).

77. Pirooznia, M. et al. High-throughput sequencing of the synaptome in major depressive disorder. Mol. Psychiatry 21, 650–655 (2016).

78. Gautier, M. & Vitalis, R. rehh: an R package to detect footprints of selection in genome-wide SNP data from haplotype structure. Bioinformatics 28, 1176–1177 (2012).

79. Sabeti, P. C. et al. Genome-wide detection and characterization of positive selection in human populations. Nature 449, 913–918 (2007).

80. Weir, B. S. & Cockerham, C. C. Estimating F-Statistics for the Analysis of Population Structure. Evolution 38, 1358–1370 (1984).

81. Grossman, S. R. et al. A composite of multiple signals distinguishes causal variants in regions of positive selection. Science 327, 883–886 (2010).

82. Grossman, S. R. et al. Identifying recent adaptations in large-scale genomic data. Cell 152, 703–713 (2013).

83. Ma, Y. et al. Properties of different selection signature statistics and a new strategy for combining them. Heredity 115, 426–436 (2015).

84. McVicker, G., Gordon, D., Davis, C. & Green, P. Widespread genomic signatures of natural selection in hominid evolution. PLoS Genet. 5, e1000471 (2009).

85. Gormley, P. et al. Meta-analysis of 375,000 individuals identifies 38 susceptibility loci for migraine. Nat. Genet. 48, 856–866 (2016).

86. Wellcome Trust Case Control Consortium et al. Bayesian refinement of association signals for 14 loci in 3 common diseases. Nat. Genet. 44, 1294–1301 (2012).

87. Lee, S. H., Goddard, M. E., Wray, N. R. & Visscher, P. M. A better coefficient of determination for genetic profile analysis. Genet. Epidemiol. 36, 214–224 (2012).

